# Condition-Specific Mapping of Operons (COSMO) using dynamic and static genome data

**DOI:** 10.1101/2022.06.14.496048

**Authors:** Tracey Calvert-Joshua, Hocine Bendou, Peter van Heusden, Melanie Grobbelaar, Rob Warren, Alan Christoffels

## Abstract

An operon is a set of adjacent genes which are transcribed into a single messenger RNA. Operons allow prokaryotes to efficiently circumvent environmental stresses. It is estimated that about 60% of the *Mycobacterium tuberculosis* genome is arranged into operons, which makes them interesting drug targets in the face of emerging drug resistance. We therefore developed COSMO - a tool for operon prediction in *M. tuberculosis* using RNA-seq data. We analyzed four algorithmic parameters and benchmarked COSMO against two top performing operon predictors. COSMO outperformed both predictors in its accuracy and in its ability to distinguish operons activated under distinct conditions.

**Author Summary:** Operons may be important drug targets for the development of effective anti-microbials to combat the emerging, global drug resistance challenge. However, there is a shortage of known *Mycobacterium tuberculosis (Mtb)* operons. This is exacerbated by the fact that current operon predictors are not optimized for the unique genome of Mtb. COSMO removes the limitations imposed by using the constraints of a specific organism’s genome and exploits RNA-seq data instead. This allows COSMO to more accurately predict full-length operons in Mtb, and it also avails COSMO to other microorganisms for the same purpose.

## 1 Introduction

Previous studies have shown that approximately 60% of the *Mycobacterium tuberculosis* (*Mtb*) genome may be arranged into operons [1]. An operon is a set of neighbouring genes that are co-transcribed as a single messenger ribonucleic acid (mRNA) [2]. These coregulated genes may not always be functionally related and may not always retain the same length [3]. Under different conditions, an existing operon may be modified by the addition or removal of one or several genes. Similarly, completely new operons may form, and old ones may be destroyed over time; demonstrating that operons are highly dynamic in their ability to evolve over both long and short periods of time. These changes often drastically alter the gene expression, and therefore also the phenotype of a species [2, 4]

However, if most genes in *Mtb* do not operate independently, then directing our anti-tubercular arsenal at individual genes may not be the most effective method of drug targeting. We need to be able to have an overview or network-level vantage-point to target larger segments of the genome more efficiently. In prokaryotes, operon structures often form in response to environmental stresses to aid these microbes to respond rapidly and efficiently in a bid to overcome adversity [5].

Fortunately, with the increasing availability of expression data in the form of RNA sequencing (RNA-seq), we are able to get an accurate representation of the overall gene expression within a species at any given moment [6]. RNA-seq is a more attractive approach than traditional platforms such as microarray, due to its wider dynamic range, ability to predict more differentially expressed genes (DEGs), and its ability to analyze a transcriptome at a single nucleotide level [7]. With an overview of a microorganism’s gene expression, we may be able to see which neighbouring genes are active and coregulated under orchestrated conditions. This data may allow us to predict operons for prokaryotes, rather than always resorting to costly and laborious experimental procedures.

Currently there are a plethora of operon predictors for prokaryotes. Tjaden (2020), who developed Rockhopper – one of the operon predictors which we benchmarked our algorithm against, tested his operon predictor on ten commonly studied microorganisms. He demonstrated that even though experimental methods may be precise and provide strong evidence, many computational tools, such as Rockhopper, can now identify operon gene pairs with predictive accuracies that exceed 90%. Several operon predictors use genomic features such as pathway analysis, sequence homology, gene ontologies and intergenic region (IGR) distance [9, 10]. Others steer away from being bound by existing genomic data and instead use statistical models to do their predictions [11]. However, to the best of our knowledge none of the existing operon predictors, except REMap, were optimized for the *Mtb* genome. The genome of *Mtb* has been shown to have some significant differences in its transcriptional preferences, such as the use of alternative sigma factors and differences in the -35 binding domains [12]. In addition, 26% of genes produce leaderless transcripts. This was especially evident in strains under a stress model [13]. REMap also showed that longer IGR lengths are common between genes of *Mtb* operons, despite this being very uncommon in other prokaryotes. Short IGR lengths are often used as the most significant feature to identify operons in existing operon predictors [11,14–17]. These all indicate that *Mtb* has unique ways of regulating its transcription, which needs to be accounted for during algorithm design.

Since the REMap algorithm considered these *Mtb*-inherent characteristics, we’ve used the foundation that REMap laid, but improved upon a few areas of their design. We’ve developed an algorithm called ‘Condition Specific Mapping of Operons’ (COSMO), which uses our existing knowledge of operons and the structural annotation of the *Mtb* genome – which we call its static data. It also uses RNA-seq to get a snapshot of the gene expression profile observed when the *Mtb* is exposed- and unexposed to rifampicin (RIF) – which we call its dynamic data. These combined data sets are leveraged by COSMO to evaluate how operons may evolve in response to RIF stress.

## 2 Results

### 2.1 Defining optimal parameters

#### 2.1.1 CDS, IGR and UTR coverages of real operons versus fake operons

We used the wiggle file to plot the **raw** coverages of the individual bases, to observe whether there was consistency in the expression patterns across genomic regions, for the 49 real operons, which did not exist in the 49 fake operons. Unsurprisingly, as displayed in Fig 1**A**, the coding sequences (CDSs) (**blue**) and their adjacent IGR coverages (**red**), showed no correlation in *fake* operons. In contrast, Fig 1**B** shows that the CDSs of *real* operons generally showed a correlation in expression levels with their adjacent CDSs, as well as with their intervening IGRs. The untranslated regions’ (UTRs) expression levels were no different between real and fake operons or single genes (plus and the minus strand: p = 0.33 and p = 0.13 respectively). The UTRs were later compared to the IGRs in **Section 2.1.4**. The UTRs served as controls to show that although the IGRs are also noncoding regions like the UTRs, within real operons IGRs are preferentially regulated and the UTRs are not, and therefore significant.

**Fig 1:**
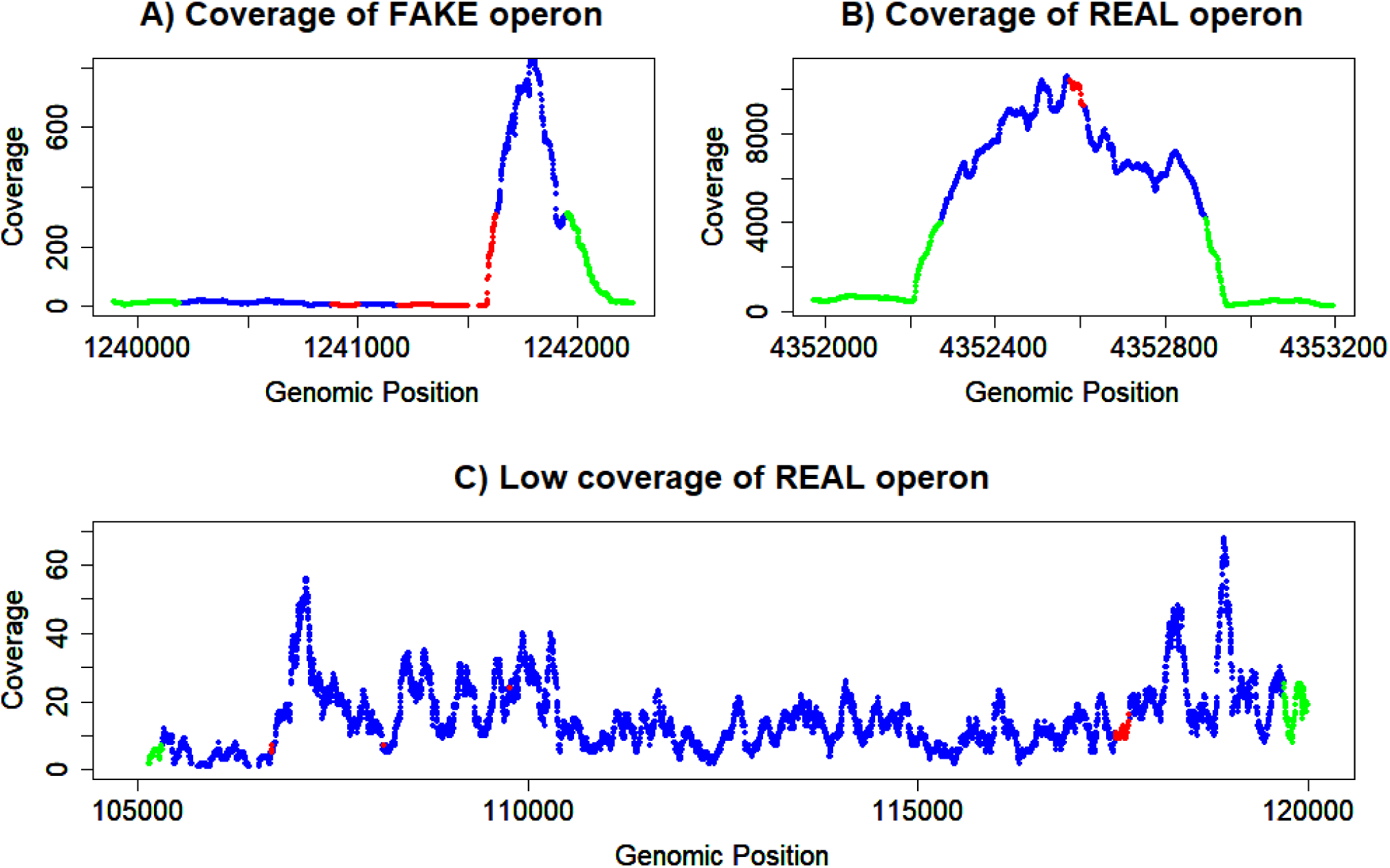
Gene expression (coverage) of CDS and IGRs for real and fake operons (Plus strand). In this graph, UTR coverages are shown in green. CDS coverages are in blue and IGR coverages are in red. **A)** There was no relationship between the CDSs and the IGRs of fake operons. In fact, in many fake operons, neighbouring CDSs were not even transcribed. **B)** The general observation for real operons, was that the expression of UTRs started to pick up *before and* trail off directly after the operon was transcribed. There seemed to be a correlation in the expression levels between the CDSs and IGRs of real operons. The UTR expression levels of fake operons were no different to those of the real operons. MWU for plus and the minus strand: p = 0.33 and p = 0.13, respectively. **C)** The expression levels of this *experimentally verified* operon demonstrates that even in real operons, some genes can have low expression levels (below 5x). Hence a strict minimum cutoff may not be feasible. Allowing the user to define the cutoff is more suitable.

#### 2.1.2 CDS coverage cutoff

The difference in expression levels between the CDSs of real versus fake operons were not statistically significant, for both the plus- and minus-strand (Mann-Whitney U test [MWU]: p = 0.22 and p = 0.65, respectively). This suggests that CDSs that make up operons are not necessarily targeted for upregulation, any more than independent CDSs (or single genes). Some of the CDSs of real operons were also expressed at very low levels - some were even below 5x coverage, as shown Fig 1**C**. Therefore, setting a high static CDS cutoff as a predictive feature, could cause the algorithm to bypass lowly-expressed or deliberately downregulated operons. The better solution might be to determine if there was a correlation of expression between adjacent CDSs of the real operons that does not exist within fake operons.

A fixed minimum value for the CDS coverage was therefore excluded as a static feature, but rather implemented as the *first user-defined parameter* in the algorithm. This is discussed later in **Section 2.2**

#### 2.1.3 Fold difference between adjacent CDSs

As anticipated, the fold differences (FD) of adjacent CDSs were more tightly regulated with respect to each other (**FD CDSs**) when they formed part of an operon, as displayed in **Fig 2A**. The maximum FDs between CDSs of real operons, were generally lower than those of fake operons (p = 0.0007). Adjacent CDSs usually adhered to a maximum FD of 5x-7x. This threshold existed not just for a CDS and its immediately adjacent CDSs, but between all the CDSs that constituted that operon - even if the operon was up to 14 CDSs long.

**Fig 2:**
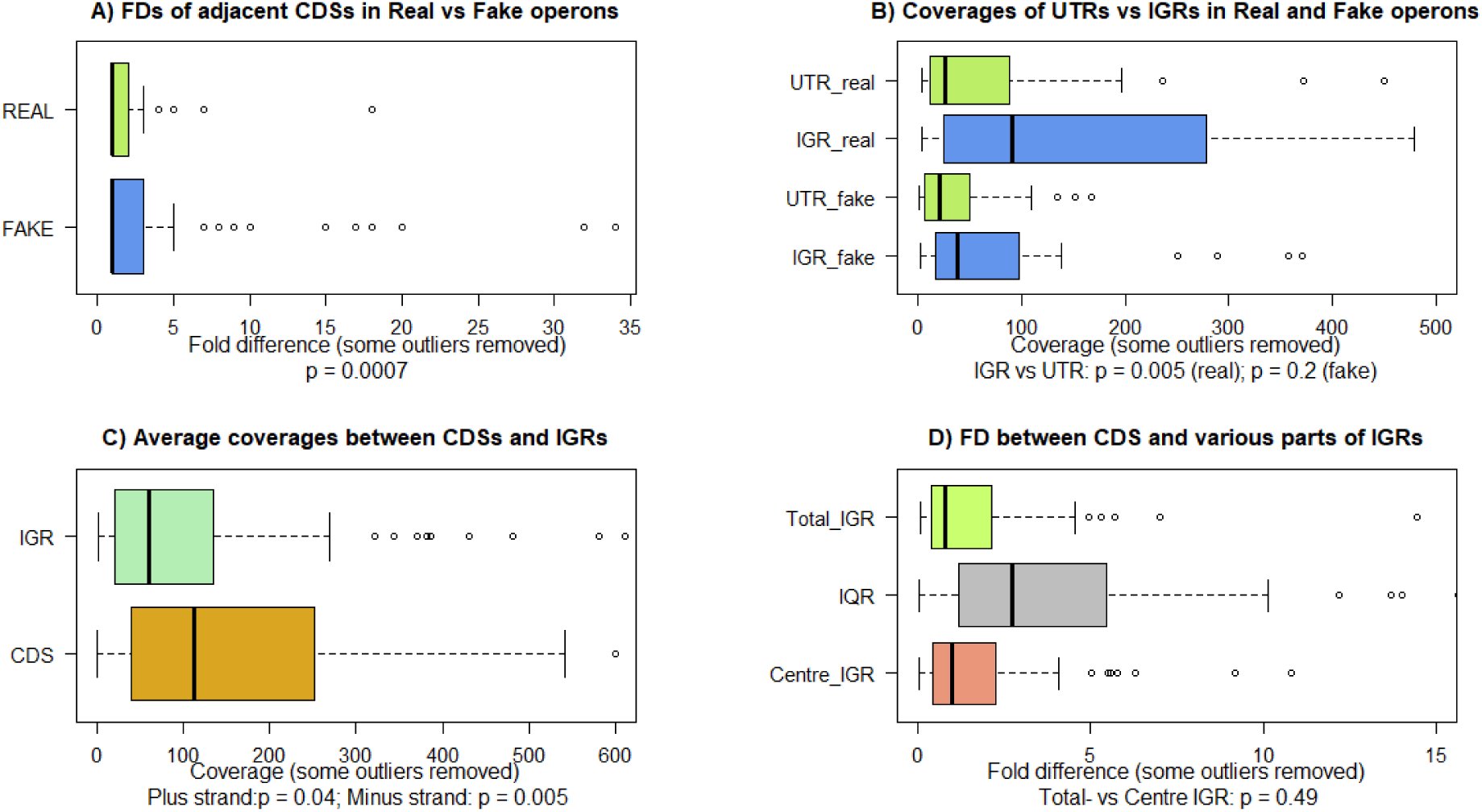
Comparison of the FDs and average coverages of the genomic regions (CDSs, IGRs and UTRs) being analyzed in real versus fake operons. **A)** The average coverages of adjacent CDSs of real operons were compared to the CDSs of fake operons. For the fake operons, some extremely large data points were removed, for a better view of the box plot. The fold FDs of adjacent CDSs in real operons usually remained within 5x-7x of each other and were also determined to be statistically significantly lower than those of fake operons (p = 0.0007). In contrast, the FDs of adjacent CDSs of fake operons often exceeded 10x. **B)** The coverages of IGRs in fake operons showed no significant differences in expression levels compared to the UTRs of fake operons (p = 0.2). In contrast, the expression levels of IGRs in real operons were more upregulated than that of the UTRs (p = 0.005). **C)** The CDS coverages of real operons were on average double that of their intervening IGRs (plus: p = 0.04 and minus: p = 0.005). **D)** The FDs of the interquartile range (IQR), the total length of the IGR and the centre of the IGR were compared to those of their flanking CDSs. Although the FDs of both the total IGR length and the centre of the IGR generally remained below 5x, the centre was chosen as the parameter for IGR coverage, since it had far fewer outliers. Some outliers were removed for better visualization of the box plots.

In contrast, the FDs of adjacent CDSs within fake operons had a larger spread, many of which also frequently exceeded 10x, or even surpassed 20x (some outliers were removed). There was however one outlier for the real operons. The FD between genes Rv3418c and Rv3419c, of the operon Rv3417c-Rv3423c, exceeded 30x. However, previous literature demonstrated that under certain stresses, this operon, could be split into two operons, namely: Rv3417c-Rv3418c and Rv3419c-Rv3423c. Thus, operon Rv3417c-Rv3418c, also known as groEL1-groES, is often expressed as an independent bicistronic operon, with the CDS Rv3418c showing evidence of gross upregulation in two experimental studies [18–20]. This was in alignment with our analysis. As a result of this exception in our already small test set, and because some operons may be better predicted with a slightly lower (or even a higher FD), we decided that we would also not restrict this value to a static maximum cutoff. Therefore, as the *second parameter* of the algorithm, users may choose their own maximum FD cutoff for adjacent CDSs. Although we do advise to keep this value to a maximum of 7x. A default FD of 5x was built into COSMO if the user does not provide their own cutoff.

#### 2.1.4 Minimum IGR expression cutoff

Regarding the IGRs, Fig 2**B** shows that in real operons, the coverages of IGRs were more *upregulated* than that of the UTRs (p = 0.005). As expected, when we similarly compared the coverages of the IGRs and UTRs for the fake operons, there was no statistical significance in their expression levels (p = 0.2).

This is in accord with our previous observations, which showed that while the CDSs and IGRs of real operons were tightly regulated - possibly by the same regulator - the UTRs are not. Additionally, the IGR coverages of real operons were also not the same as the CDSs but were on average 50% lower than the coverages of their flanking CDSs, as depicted in **Fig 2C** (p = 0.04 and p = 0.005; plus- and minus-strand, respectively). Therefore, the IGR coverage is not just a significant parameter when contrasted with the UTRs, but it should be and independent parameter relative to their CDSs. IGR coverage was therefore included as the *third user-defined parameter* in COSMO.

We then wanted to obtain a minimum cutoff for an IGR to be considered expressed. However, just as with the CDSs, the IGRs of real operons were generally not more up- or downregulated compared to individual IGRs of fake operons. The MWU test showed that the outcome was inconclusive. There was a statistically significant difference for the minus strand (p = 0.01), but not for the plus strand (p = 0.14). However, even though there may not be a defined minimum cutoff for the IGR coverage, from **Fig 1B**, we saw that in real operons, the IGRs show a correlation of expression levels with their adjacent CDSs. In contrast, **Fig 1A** showed that the expression levels of IGRs and adjacent CDSs, behave haphazardly in fake operons. This suggests that just as with the CDSs of real operons, the IGRs may stay within a maximum FD to their adjacent CDSs.

#### 2.1.5 Fold difference between IGR and adjacent CDSs

As depicted in Fig 2**D**, the possibility of using FDs between the interquartile ranges (IQRs) of the IGRs and their flanking CDSs were immediately discarded, because the spread of the data points representing the FDs, was too large and too random. Next, the FDs for the total IGR length and for that of the centre of the IGR were analysed. The boxplot shows that either one of the two may have been used as a source to calculate the FDs between the IGR and its flanking CDSs, because they did not perform very different (p = 0.49). However, when the total IGR lengths were used, there were more outliers. Hence, the algorithm utilizes the centre of the IGR to establish the FD between an IGR and its adjacent CDSs. The maximum FD for an IGR and its flanking CDSs was therefore also included as the *fourth user-defined parameter* of the algorithm.

#### 2.1.6 Intergenic distance

We then evaluated if IGR distance is an appropriate feature/parameter for *Mtb* operon prediction. As depicted in the density plot in Fig 3**A**, the peaks confirmed that most CDSs overlapped in real operons (had no IGR). Still, Fig 3**B** reveals that even when using 50bp, as opposed to the 20bp usually considered, the lengths of 33% of IGRs on the minus strand and 16% of IGR on the plus strand exceeded that which is normally observed in other prokaryotes. Our results therefore supported the findings of REMap, that a maximum IGR length may not be a useful feature for operon prediction in *Mtb*. We found that the length of the operon also had no impact on the length of the IGRs. Meaning, IGR lengths longer than 20bp were observed as frequently in operons containing two CDSs as they were in operons that were 14 to 15 CDSs in length. However, there was a preference of location for longer IGR lengths. In 44% of cases, excessive IGR lengths were between the first two CDSs of an operon. It should be noted that eight of these long IGRs were within operons containing just two CDSs. Hence, in the cases of bi-cistronic operons, we will miss entire operons if we filter and exclude CDSs based on IGR length. In fact, in many instances, these long IGRs were also between all the CDSs of longer operons. We calculated that just under a 1/3^rd^ (26%) of our real operons would not be predicted if IGR length is used as a feature. This parameter was therefore excluded from this algorithm.

**Fig 3:**
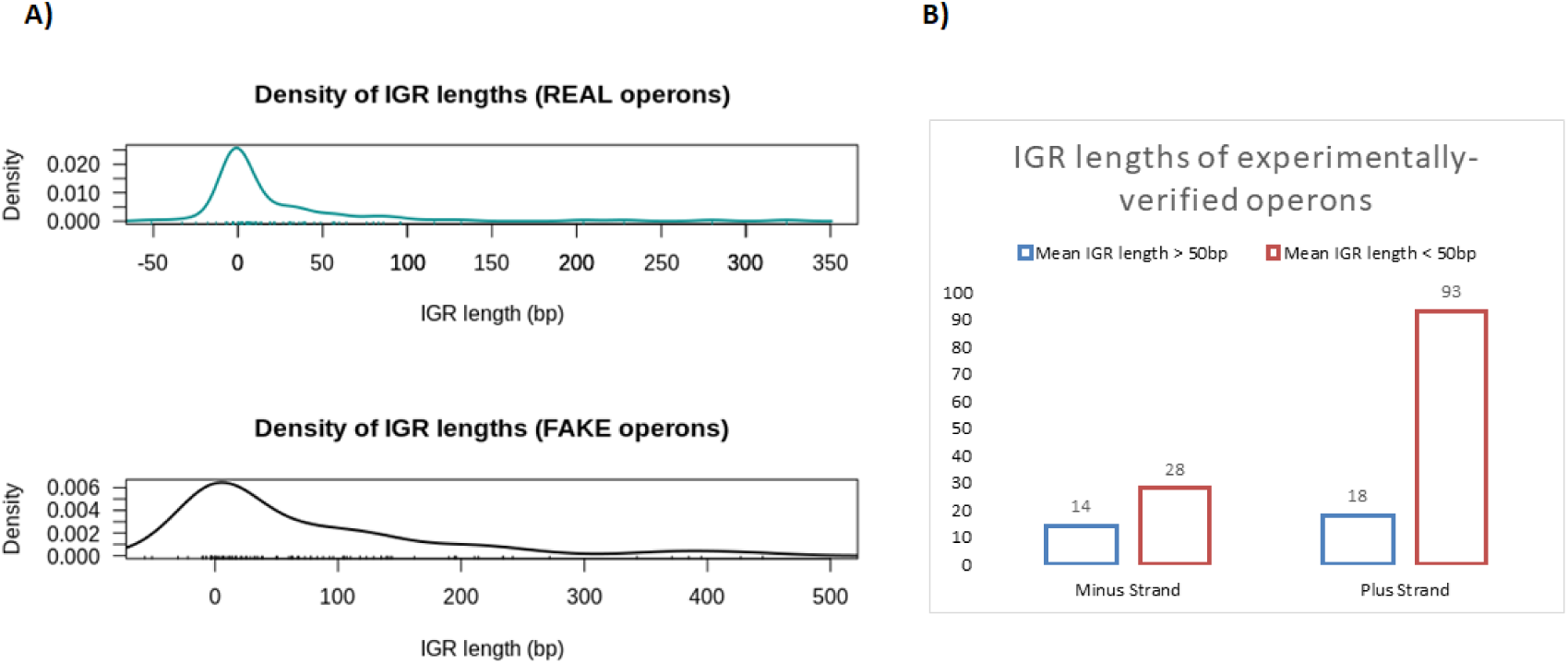
Comparison of Intergenic distance between real and fake operons. **A)** Although the IGR lengths peaked around 0 nucleotides, there were still far too many IGR lengths that were longer than what is usually observed in operons of other prokaryotes. **B)** About 1/3^rd^ IGRs on the minus strand (33%) and 16% of IGR on the plus strand exceeded 50bp. Limiting the operon predictions by IGR length would result in the exclusion of 1/3^rd^ of verified operons. This parameter was therefore excluded from the algorithm.

### 2.2 Algorithm validation

#### 2.2.1 Multiple Linear Regression (MLR) Analysis

We anticipated that the FD between adjacent CDSs would have the greatest effect on the outcome. Surprisingly, the MLR analysis showed that the greatest impact on the outcome variable was the maximum FD between the IGR and its flanking CDSs (coefficient estimate = -0.37). Meaning that each time this parameter is **decreased** by just 1 unit, the total number of operons predicted, increases by 0.37 percent. Naturally, this number is not very high, because our verified TP list is small, so there are less operons to catch. The next most significant parameter was the min CDS coverage, followed by the min IGR cutoff and at the very least, the max FD between adjacent CDSs. The MLR showed that all four predictor variables were highly statistically significant (< 3.8 x 10^-16^). As expected though, despite their significance, these variables accounted for 40% of the variability (adjusted coefficient of determination [R^2^] = 0.4). We suspect that the other variable (splitting putative operons at the point where correlation breaks between distant adjacent CDSs), discussed in **Section 5.2**, may have a large impact on the outcome, but this could unfortunately not be switched on and off. Despite being routinely used, several authors have argued against using R^2^ as a strict predictive measure of model’s performance. They argued that R^2^ may be a biased, insufficient and misleading measure of predictive accuracy, and that Root Mean Squared Error (RMSE) may give a much better indication of the accuracy of a model [21, 22]. We therefore measured the RMSE and the results showed that the error rate was definitely reduced in the final model (2.6), compared to the baseline model (3.4). Similarly, the mean absolute error (MAE) dropped from 2.5 to 2.1 in the final model, showing that the decision tree performs better when these four parameters were used, than if they were excluded. The pruned tree model (which combats overfitting of data) again calculated no default cutoff for a max FD between adjacent CDSs, but it computed that the default cutoffs that can be utilized to correctly predict >=37/50 experimentally verified operons (EVOs) for most strains, would be achieved if we restrict the:

i) **5.5 <** FD between CDS and flanking IGRs **<=13**,
ii) min CDS coverage **<= 7.5** and
iii) min IGR coverage **<= 6.5**.

These values confirmed our previous observations and were therefore considered for the default parameters of COSMO.

#### 2.2.2 Comparison to existing algorithms

Finally, we compared the total full-length operons called by COSMO to REMap and Rockhopper. We settled on using the 10x cutoffs for REMap as per their publication, because it performed better than their algorithm’s default setting of 20x. As per the REMap algorithm, this cutoff applies to both the CDS and IGR coverages.

COSMO with its four parameters as input (**Control**: min CDS = 1x; min IGR = 4x; max FD of IGR-vs-CDS = 6x; max FD adjacent CDSs = 7x. **Experimental:** min CDS = 2x; min IGR = 1x; max FD IGR-vs-CDS = 5x; max FD adjacent CDSs = 5x ) was able to accurately predict more operons under both the control and experimental conditions (52% and 50%, respectively) than REMap (44% and 46%, respectively) and Rockhopper (48% in total), as shown in **Table 1**. This is significant because most existing algorithms do not generate conditions-specific operons. Rockhopper for example, has only a total value, because it predicts operons based on differential expression and Rockhopper also does not allow user-defined cutoffs like REMap and COSMO. When the control and treated samples are submitted independently for operon predictions, Rockhopper generated identical reports.

**Table 1:**
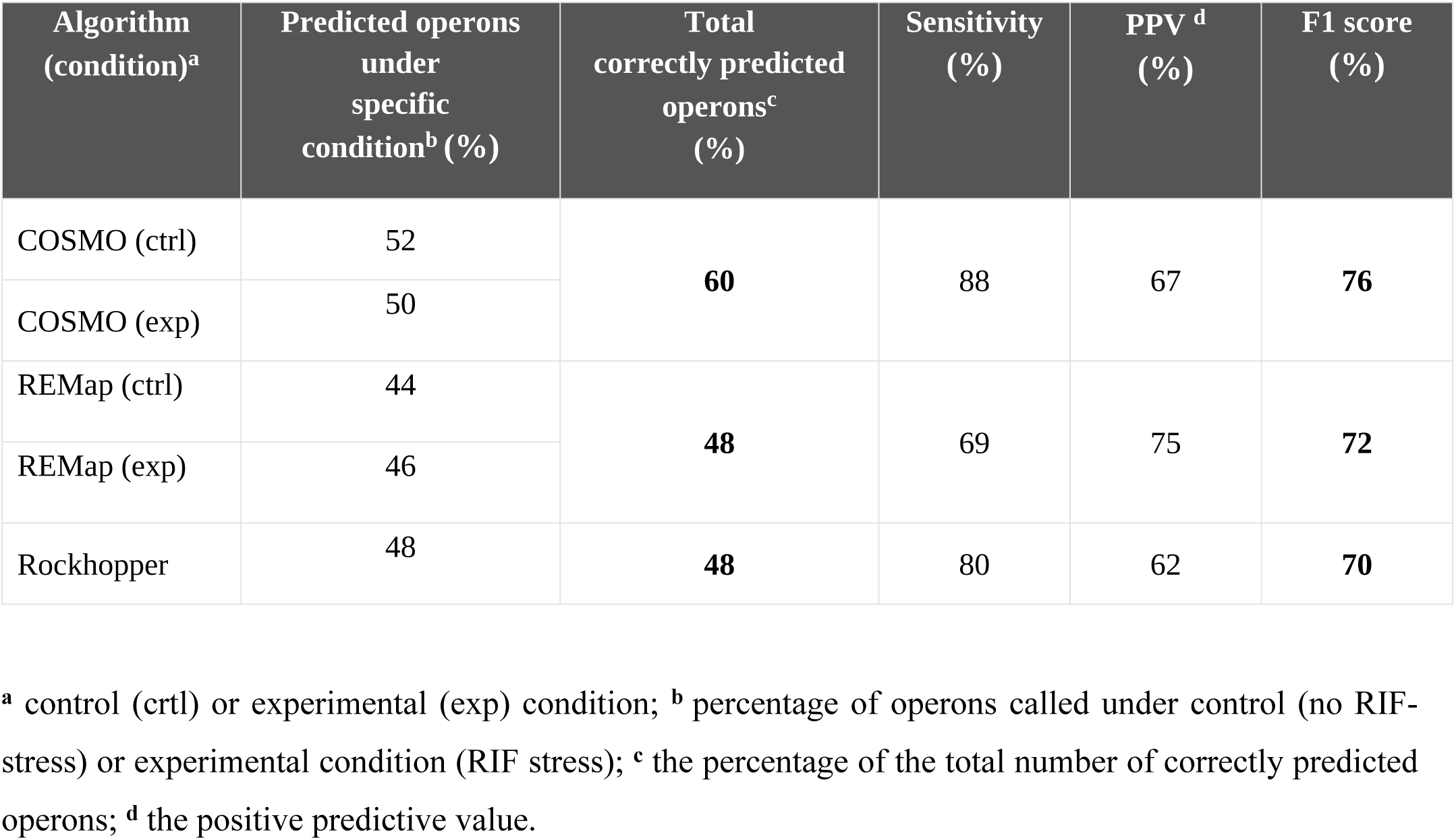
Comparison of the performance of the three operon prediction tools: COSMO, REMap and Rockhopper.

| Algorithm (condition)<sup>a</sup> | Predicted operons under specific condition<sup>b</sup> (%) | Total correctly predicted operons<sup>c</sup> (%) | Sensitivity (%) | PPV <sup>d</sup> (%) | F1 score (%) |
| --- | --- | --- | --- | --- | --- |
| COSMO (ctrl) | 52 | **60** | 88 | 67 | **76** |
| COSMO (exp) | 50 |  |  |  |  |
| REMap (ctrl) | 44 | **48** | 69 | 75 | **72** |
| REMap (exp) | 46 |  |  |  |  |
| Rockhopper | 48 | **48** | 80 | 62 | **70** |

<sup>a</sup> control (ctrl) or experimental (exp) condition; <sup>b</sup> percentage of operons called under control (no RIF-stress) or experimental condition (RIF stress); <sup>c</sup> the percentage of the total number of correctly predicted operons; <sup>d</sup> the positive predictive value.

Moreover, when the number of operons predicted under both conditions were combined, COSMO’s <u>total</u> predicted operons (60%) also exceeded that of REMap (48%) and Rockhopper (48%). COSMO was also the frontrunner with regards to sensitivity (88%), compared to REMap (69%) and Rockhopper (80%). This however came with the usual trade-off in precision where REMap performed better (75%) than COSMO (67%) and Rockhopper (62%). However, overall, COSMO was not only correctly identifying more operons, but it was also doing it more accurately, with F1 scores for COSMO, REMAP and Rockhopper at 76%, 72% and 70%, respectively.

We are aware that the total number of operons caught may still seem low. In many other studies where operons were predicted, the performance metrics are often over 80% or 90% for sensitivity, specificity, F1 and accuracy scores [23–26]. However, in these studies, operons are usually split into gene pairs. This obviously leads to a much bigger true positives (TPs) test set, allowing for more correct predictions to be made. With COSMO, if an operon consisting of 5 genes is split after CDS 2, our algorithm computes it as unpredicted, or a FN, whereas in in other studies, since only one gene pair was not predicted, it will be counted as 3 TPs + 1 FN.

#### 2.2.3 Unique predictions

In terms of unique predictions from the list of EVOs, seven operons were predicted only by COSMO, as displayed in Fig 4**A**, and three operons were predicted by Rockhopper. REMap was also able to predict one operon from literature that was not predicted by the other two algorithms. This is discussed in greater detail in **Section 2.2.5**.

**Fig 4:**
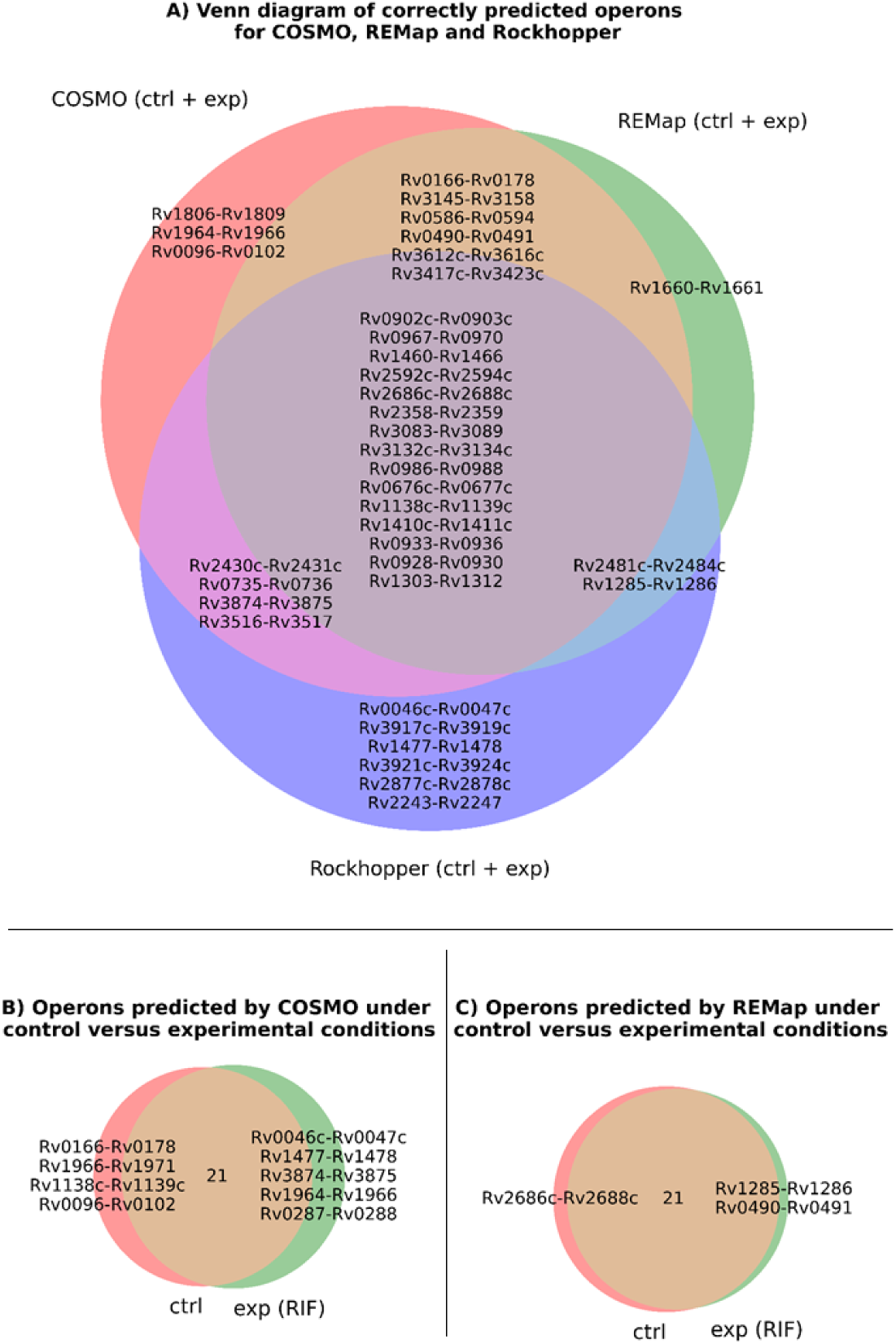
Comparison of operons predicted by COSMO, Rockhopper and REMap. **A)** The intersection of operons from COSMO, Rockhopper and REMap. Only one operon was uniquely predicted by REMap, while three operons were uniquely predicted by Rockhopper. Seven operons were predicted solely by COSMO. **B)** COSMO also predicted four operons as only expressed under control conditions, while five operons were predicted only under RIF stress. **C)** REMap was also able to distinguish two operons that were predicted under control conditions from the one specific to RIF stress. Rockhopper is not shown, because the algorithm only predicted combined differentially expressed operons.

#### 2.2.4 Condition-dependent mapping

Fig 4**B**, illustrates the ability of COSMO to distinguish between operons predicted under control conditions from those predicted when under stress (RIF treatment). Four operons were predicted solely under control conditions, while five operons were predicted as active only under RIF stress. As shown in Fig 4**C**, although REMap predicted less operons than COSMO, it also demonstrated the ability to classify operons in a condition-specific manner - a distinction Rockhopper was not able to make. The Venn diagrams were created using ‘*matplotlib-venn’* 0.11.6 [27] in Jupyter Notebook 6.0.3 [28].

#### 2.2.5 Operons not predicted by COSMO

We investigated the operons not predicted by each algorithm, to determine whether the operons were either shorter than those reported in literature (FNs), or longer (FPs). **S1 Table 1** shows the 50 EVOs and their prediction calls across the three algorithms. Only 8% (n = 4) of the operons incorrectly predicted by COSMO were shorter than those in literature - or could be considered as FNs; in contrast to REMap and Rockhopper for which 20% (n = 10) and 29% (n = 14) of operons were FNs, respectively. Most of the operons not found by COSMO were predicted to be slightly longer than those in the literature and none of the operons were completely unexpressed (zero genes expressed). On the contrary, REMap and Rockhopper called all the genes of some operons as unexpressed (n = 6 and n = 5, respectively). Although DOOR 2.0 no longer predicts operons, we compared our results to a list of operons already predicted by DOOR 2.0 for *Mtb* (not shown). COSMO also outperformed DOOR, by predicting nearly twice as many operons as DOOR, which correctly predicted only 16 of the total operons (33%).

One especially interesting feature of COSMO is that it is able to predict operons with CDSs that are expressed at very low levels. As briefly discussed in **Section 2.2.2**, the expression levels of ∼5x, for both the CDS and IGR, resulted in the highest number of total predicted operons for most isolates. This is because COSMO is able to bypass low expression levels, while rather taking advantage of a maximum FDs between CDSs and between an IGR and its flanking CDSs. This is also one of the reasons why REMap and Rockhopper were not able to predict some operons from literature. One example of this is the operon Rv3516-Rv3517, which was caught by COSMO, but not by REMap or Rockhopper, since its expression levels were very low. Therefore, if the expression of an operon is deemed detrimental by *Mtb* for its virulence or survival or it’s biologically redundant, and it deliberately downregulates the expression of this operon, COSMO would still be able to predict those operons and record the downregulated expression.

#### 2.2.6 Composition of predicted operons

Lastly, COSMO’s potential to predict novel operons was also analyzed. Under control- and RIF stress conditions, the number of putative operons and the number of CDSs forming part of putative operons were on average not notably different across isolates. Table 2 shows the results for one particular isolate. Under control conditions, COSMO predicted 75% genes of the total 4109 protein coding genes in *Mtb* to be constituents of operons (n = 941 operons). We predicted that 23% were expressed as single genes and 2% were single unexpressed genes. When under RIF stress, the genome seems to generally be more active, as there were fewer unexpressed genes. However, this was not always the case. For a few isolates, more single genes were predicted as expressed under control conditions. Therefore, there were only three consistent characteristics when the predictions for all isolates were analyzed. The first was that COSMO always predicted a higher number of operons under RIF stress compared to control conditions. The second was that operons were usually predicted to be shorter in length under RIF stress. The largest operon predicted was under the control conditions, consisting of 17 CDSs. COSMO was therefore in agreement with the results that REMap and Rockhopper previously showed, in that at any given point in time, the larger proportion of *Mtb* genes/CDSs, are not operating independently, but they are instead predicted to be constituents of operons. COSMO predicted that up to 75% of the *Mtb* genome may constitute operons – whether it was control or RIF-stress conditions. REMap reported this number to be just under 60%, while Rockhopper predicted this as 62% in other prokaryotes.

**Table 2:**
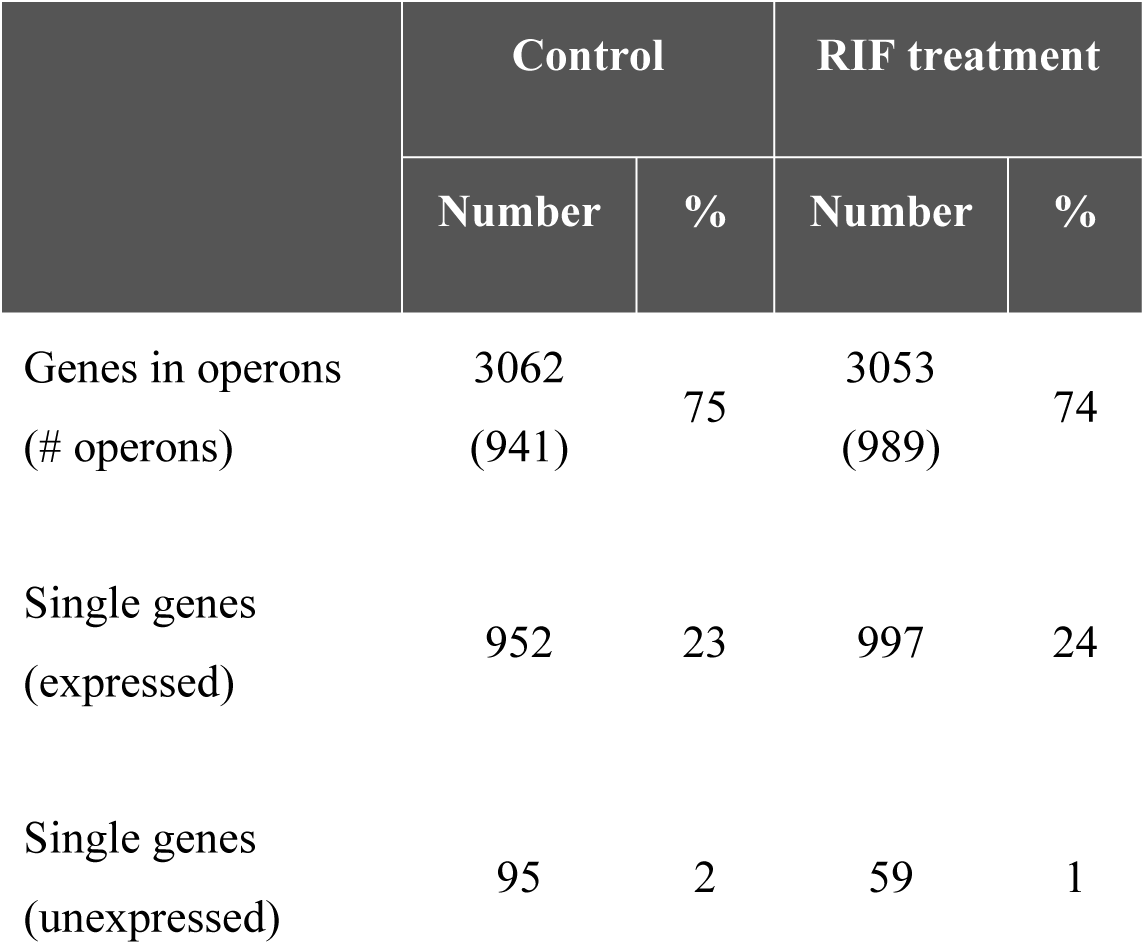
Percentage of the Mtb genome predicted as operons and single genes. COSMO predicted that approximately 75% of the *Mtb* genes form part of operons. Less than a ¼ of *Mtb*’s total protein coding genes are predicted to be expressed as single gene entities and approximately 1-2% of single genes are predicted as unexpressed at any given time point.

|  | Control |  | RIF treatment |  |
| --- | --- | --- | --- | --- |
|  | Number | % | Number | % |
| Genes in operons<br>(# operons) | 3062<br>(941) | 75 | 3053<br>(989) | 74 |
| Single genes<br>(expressed) | 952 | 23 | 997 | 24 |
| Single genes<br>(unexpressed) | 95 | 2 | 59 | 1 |

##### Availability of data and materials

• The data discussed in this publication have been deposited in NCBI’s Gene Expression Omnibus (Edgar et al., 2002) and are accessible through GEO Series accession number **GSE203032** (samples GSM6152783 to GSM6152802): https://www.ncbi.nlm.nih.gov/geo/query/acc.cgi?acc=GSE203032 The code and examples of input and output files for testing and validation, are available at the project home page at https://github.com/SANBI-SA/COSMO The archived version is at https://zenodo.org/record/6483309

## 3 Discussion

COSMO takes in RNA sequence reads (.bam files) together with a GTF file and identifies the operons active under varied experimental conditions. For this study, we used RIF-treatment as our experimental condition and no RIF treatment as the control. The user is able to provide four separate parameters as cutoffs. Although other RNA-seq based algorithms also allow users to define an expression level cutoff, to the best of our knowledge this is the first algorithm predictor that allows the user to provide a separate input for IGR coverage and CDS coverage. It is also the first operon predictor where users can define the maximum FD between adjacent CDSs as well as the maximum FD between an IGR and its flanking CDSs. Additionally, COSMO not only considers correlation of expression between directly adjacent genes, but between all CDSs within a predicted operon.

Allowing the user to provide the FDs is especially valuable. These parameters can take pre-eminence over just considering the min CDS or IGR expression levels as predictive indicators. This tolerates expression levels of CDSs and IGRs that are very low, which is crucial for identifying operons that may have been downregulated by the bacteria in response to stress.

COSMO can distinguish between operons predicted under control conditions and operons predicted under experimental conditions. Being able to match which operons are expressed under each condition is valuable, because we may observe which pool of genes are specifically expressed under one condition, while being deemed either detrimental or at the very least, redundant under another condition. This capability may eventually allow us to better understand how *Mtb* is able to circumvent and thrive under stress conditions by tailoring a condition-specific pool of genes within the boundaries of an operon, as well as which operons are never altered, but may be considered as “housekeeping operons”.

COSMO was able to predict all experimentally validated operons from literature, with 60% of the operons being exact matches to those obtained from literature, while Rockhopper and REMap predicted 48% each. COSMO also obtained a greater F1 score (76%), compared to REMap (72%) and Rockhopper (70%). With regards to those that were not exact matches; some operons were slightly shorter than in literature. However, most operons that were not identical to those from literature, were predicted to be slightly longer. On the contrary, REMap and Rockhopper not only predicted more shorter operons, but they also predicted entire operons as unexpressed. Although matching only 60% of operons from literature correctly may seem rather low, it would be more alarming if we predicted everything or if we predicted close to 100% of operons. This would contradict our understanding that different operons are expressed under different environmental conditions. For the purpose of this study, we have only tested COSMO on isolates under control versus RIF stress conditions. This was because the main objective was just to evaluate COSMO against existing operon-predicting algorithms.

COSMO also predicted that up to 75% of the genome are organized into operons, which may indicate that *Mtb* has a heavy reliance on forming operons as a means of regulating its genome. We also predicted that control conditions may favour longer, but fewer operons. On the contrary, under RIF stress, shorter operons were predicted, but they were more frequent. This may suggest that under RIF-stress, *Mtb* activates operons on a *ad hoc* basis, in order to swiftly and efficiently handle the adversity it faces at that specific time. The significance of this, is that *Mtb,* the species responsible for causing Tuberculosis, is known for its impressive ability to evade and survive within their hosts [29]. This pathogen, which is responsible for more deaths than any other infectious agent, worldwide [30], has co-evolved with its hosts over several millennia and has continuously outsmarted the myriad of drugs that were carefully designed to disrupt its virulence at a gene or single nucleotide polymorphism (SNP) level [31, 32]. If *Mtb* favours operons under stress conditions, then it may make more sense to study its evasive tactics in the context of operons, rather than by looking at mutations or differential expression of individual genes – which is how it was traditionally done. However, further analyses would have to be carried out to determine whether creating shorter operons allows it to have a tighter control over gene regulation or whether it has an alternative purpose. One of our current analyses involves taking a deeper look at the functions of the genes in the operons that are changing and dysregulated under each experimental condition.

### Limitations

One of the limitations we had in this study was the very small list of validated operons. Once this list becomes more populated, we may be able to evaluate the algorithm’s accuracy using more traditional methods such as sensitivity- and specificity receiver operating characteristic (ROC) curves. We are hoping that once this algorithm is tested across a variety of lineages and experimental conditions, we may be able to detect the static CDSs of operons. This may aid us in further optimizing the algorithm if static operons can lead us to one or a number of consensus motifs that can be used as a feature or parameter in the algorithm design. Lastly, an experimental validation will have to ensue on carefully selected candidate operons predicted by COSMO to further gauge its performance.

This analysis will also be extended to other *Mtb* families and to *Mtb* genomes exposed to different environmental conditions. This should generate a higher number of matches to operons published in literature, since the current experimentally validated operon list we used, consisted of operons discovered from a variety of different *Mtb* lineages and from a variety of experimental conditions.

## 4 Conclusion

COSMO has demonstrated an improved capacity to identify existing operons when compared to REMap and Rockhopper. Additionally, because it does not rely on inherent *Mtb*-specific traits for operon prediction, it could also be utilized for operon predictions in other microorganisms. This will allow us to break through the current bottleneck of focusing primarily on individual genes, without understanding the regulatory mechanisms at play and potentially lead to more potent drug targets as well as a greater understanding of an organism’s genome.

## 5 Methods

### 5.1 Strains and growth conditions

Samples were obtained from two tuberculosis patients, from which we isolated nine strains belonging to the Beijing lineage. The lineage and drug resistance profile were confirmed using the *TBProfiler* tool [33] in Galaxy (https://galaxy.sanbi.ac.za/) [34]. The strains were classified according to their rifampicin minimum inhibitory concentration (MIC), with five having a high MIC (150 ug/ml) and four having a low MIC (40 ug/ml). Cultures were grown in 7H9 media until mid-log phase and exposed to RIF for 24 hours. Both the high- and low MIC strains were exposed to ¼ MIC of rifampicin. The control batches received no RIF treatment.

### 5.2 RNA extraction and sequencing

RNA extraction was carried out using the FastBlue RNA extraction kit (MP Biomedicals). Ribosomal depletion was performed, with the bacterial option as probes for hybridization of rRNA (TruSeq Total RNA). Primer design and RNA-seq were done by the Agricultural Research Council (ARC) sequencing facility using the TruSeq DNA and RNA CD Indexes (I7 and I5 adapters). RNA-seq was performed using Illumina 1.9 with FR firststrand strandedness.

### 5.3 Trimming and alignment

A FastQC [35] check in Galaxy showed that the data was of high quality (mean PHRED > 30), but the reads were nonetheless trimmed, and adapter sequences were removed using Trimmomatic V.0.38 with a PHRED = 20 and a sliding window of 4 [36]. Reads were aligned to H37Rv (NC_00096.3) using BWA-MEM 0.17.1 [37] and the quality of the bam files were checked using Samtools 1.9 [38]. Some of the bam files were converted to wiggle files for further analysis, using the *RSeQC* package in Galaxy [39].

### 5.1 The Algorithm Design

Using an RNA expression analysis with microarray data, Tjaden et al., (2002) noted that including similar expression levels for the IGR which is flanked by two adjacent operon genes, provided a much stronger signal for operon detection than just using the expression of adjacent genes/CDSs. This approach also yielded a false positive rate that was below 1%.

Although Pelly et al. (2016) used this insight and also took the complexity of the *Mtb* genome into account during their design of REMap, they used only one user-defined parameter for both the minimum CDS- and IGR coverage. However, Okuda et al. (2007) showed that the IGRs had distinct expression levels in operon gene pairs. REMap also did not consider the correlation of expression between two adjacent CDSs or between an IGR and its adjacent CDS. We opted to improve on the foundation laid by REMap by extending the algorithmic parameters with empirical evidence. With COSMO we verified **four parameters**, by evaluating a set of 49 experimentally confirmed operons and a matching simulated operon set – which we call the *fake* operons. The 49 operons were obtained from literature from both the plus- (n = 30) and the minus strand (n = 19). We created a set of 49 fake/simulated operons, using adjacent genes that were **not** previously confirmed to belong to an operon and which did not overlap with, and were not located in close proximity to verified operons. Each fake operon therefore matched its real operon counterpart by strand and by the number of genes. This true negative operon list was only used for this initial comparison between the coverages of real and simulated operons. We were always aware that they were not verified true negatives, but that they may have actual operons that have not yet been discovered. Therefore, this list was not used as a true negative (TN) set to measure the performance of the three algorithms.

Some of the parameters we aimed to include and assess for this algorithm were:

i) how many reads should be available on average for a gene or coding sequence (CDS) to be considered expressed (minimum CDS coverage)?
ii) is there a correlation between expression levels of CDSs of the same operon? (maximum FD between adjacent CDSs)
iii) Should the intergenic region (IGR) be an independent parameter and if yes, then what should be the minimum coverage to consider it expressed (minimum IGR coverage)?
iv) is there a correlation between the expression levels of IGRs and its flanking CDSs?
v) should the entire IGR be used for comparison to flanking CDSs or is the relationship between IGR and CDS more tightly regulated with certain parts of the IGR?
vi) should we use IGR length/distance as a feature?

#### 5.1.1 Defining and extracting genomic features

The coverages were extracted for each CDS, IGR and UTR, for both the real and the fake operons, using the wiggle file, as explained in **Section 5.3**. A wiggle file specifies the depth of aligned reads in a per-base format. The CDS is defined as the protein coding sequence, and for the purpose of this study, it was defined by the coordinates in the NC_00096.3 GTF file (Ensembl). The IGR was defined as the region between two adjacent CDSs on the same strand. The UTRs were taken as the regions 300 bases up- and downstream of the start- and end coordinates of operons. We used 300 bases, since this was below the maximum length of the longest IGRs in our dataset, and also the median value for long 3’ and 5’ UTRs [42, 43]. We exploited this data to compute the coverage depth, which is calculated using the number and length of reads mapped for each genomic position.

The MWU test was used to determine if there was a statistically significant difference in the average coverages and FDs, between the genomic regions of real and fake operons, using base R v3.6.1 [44].

#### 5.1.2 CDS, IGR and UTR coverages of real operons versus fake operons

First, we wanted to ascertain whether there were **observable differences** in the overall expression patterns of operons versus non-operons (fake operons). Based on the wiggle files generated from the aligned RNAseq reads, we drew plots in R studio. We hypothesized that in the real operons the coverages of adjacent CDSs and their intervening IGRs should be correlated, while they should show no correlation in the fake operons. The UTRs served as an additional control, because the UTR expression levels should technically not be under selective pressure for coregulation in both the real or fake operons, and therefore their expression levels were expected to **not** differ. In addition, by comparing the noncoding UTRs of real operons to the IGRs of that same operon, one should be able to observe that while the UTRs may be uncorrelated to the operon CDSs, the IGRs should show preferential correlation to their flanking CDSs. On the contrary, both the expression of the IGRs and UTRs should be uncorrelated to the adjacent CDSs in fake operons.

#### 5.1.3 CDS coverage cutoff

The first genomic region which we considered, was whether there should be a minimum expression level cutoff for CDSs of **real operons** (**min-CDS**), that could be set as a feature or parameter for operon prediction. If ***a*** is considered to be the start of a CDS, and ***b*** is considered to be the end of a CDS, then:

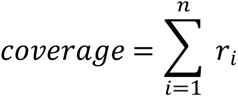

Where r_i_ is the total number of reads that mapped to the nucleotide position *i*.

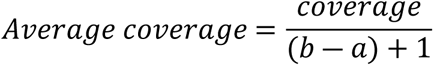

The average coverages of the IGRs and UTRs were calculated similarly, using their specific genomic positions from the GTF file.

#### 5.1.4 Fold difference between adjacent CDSs

To determine the correlation of expression levels between adjacent CDSs of operons, the FDs between adjacent CDSs (**FD CDSs**) within the same operon were calculated using a Python script, where:

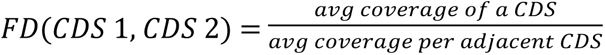

#### 5.1.5 Minimum IGR expression cutoff

We then considered the relevance of the IGRs. For the initial part of the analysis, we assessed if there was a statistically significant difference between the expression levels of IGRs versus UTRs in real operons compared to fake operons, since they are both non-coding regions. This was previously done only by observation. Then in the second part of the analysis, we tested whether we should use a minimum IGR coverage (**min-IGR**) as a user-defined parameter.

#### 5.1.6 Fold difference between IGR and adjacent CDSs

We then assessed the relationship between the IGRs and their flanking CDSs. In computing the average expression level of an IGR we need to bear in mind that expression levels increase before the transcription of a gene (i.e. before the 5’ end) and tail off after the 3’ end. For IGRs longer than four base pairs we split the IGR into four regions and computed the average expression from the middle two segments (see **S2 Fig 1**), thereby avoiding the ramp up and tail off effects mentioned above. For IGRs shorter than four base pairs we compute the average expression from the entire IGR.

We recorded the FD for an IGR and its preceding CDS separate from the FD between an IGR and its succeeding CDS, to see if there was a difference. The FD between the IGR and its flanking CDSs (**FD CDS-IGR**) was calculated as,

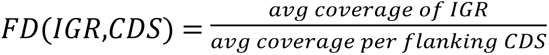

#### 5.1.7 Intergenic distance

We then compared the average IGR lengths in real operons to the average IGR lengths of fake operons. As previously stated, in many operon predictors, the intergenic distance is considered the most defining feature for accurate operon prediction. This is due to the finding that in most prokaryotes, the adjacent CDSs of operons often either overlap, or the IGR distances between adjacent CDSs are separated by fewer than 20bp of DNA. However, Pelly et al. (2016) reported that with *Mtb*, this distance was often 200 nucleotides long, but could reach up to 2.47kb in length. We calculated the lengths of IGRs of verified operons to observe if most IGRs were under 50bp. We used 50bp instead of 20bp to give some leeway to the longer IGRs. This was then used to consider it as a potential predictive feature.

### 5.2 Decision Tree

After the selected features and parameters were statistically validated, COSMO was designed using the decision-tree based classifier. The decision tree classification method was previously tested in many different operon predictors and found to produce the highest sensitivity and specificity values [14]. It takes in a BAM file, a GTF file, as well as four user-defined parameters: ***a)*** a minimum CDS cutoff, ***b)*** a minimum IGR cutoff, ***c)*** the maximum FD between an IGR and its flanking CDSs and ***d)*** the maximum FD between two adjacent CDSs.

As shown in Fig 5, COSMO starts by checking if the first CDS it encounters is expressed. That is, it checks if the average CDS coverage is equal to or above the user defined CDS cutoff **(*a*)**. If it is expressed, it then assigns it as CDS 1 of a putative operon. It then advances to CDS 2 on the same strand. If CDS 2 is expressed and it overlaps with CDS 1, it automatically gets added as CDS 2 of the operon. Should CDS 2 not overlap CDS 1, then the IGR has to be expressed. That is, the average coverage of IGR must be equal to or above the user-defined cutoff **(*b*)**. The average of the entire IGR region is considered if its length is below 4 nucleotides. In the case where the IGR between CDS 1 and CDS 2 exceeds 4 bases, the IGR is split into four parts and the average of the bases making up the two middle regions is used for further analysis. Next, the FD between this IGR coverage and each adjacent CDS must not exceed the maximum IGR to CDSs cutoff **(c)**. Finally, before adding CDS 2 to the operon, the FDs between CDS 1 and CDS 2 must not exceed the maximum CDS-to-CDS cutoff ***(d)***.

**Fig 5:**
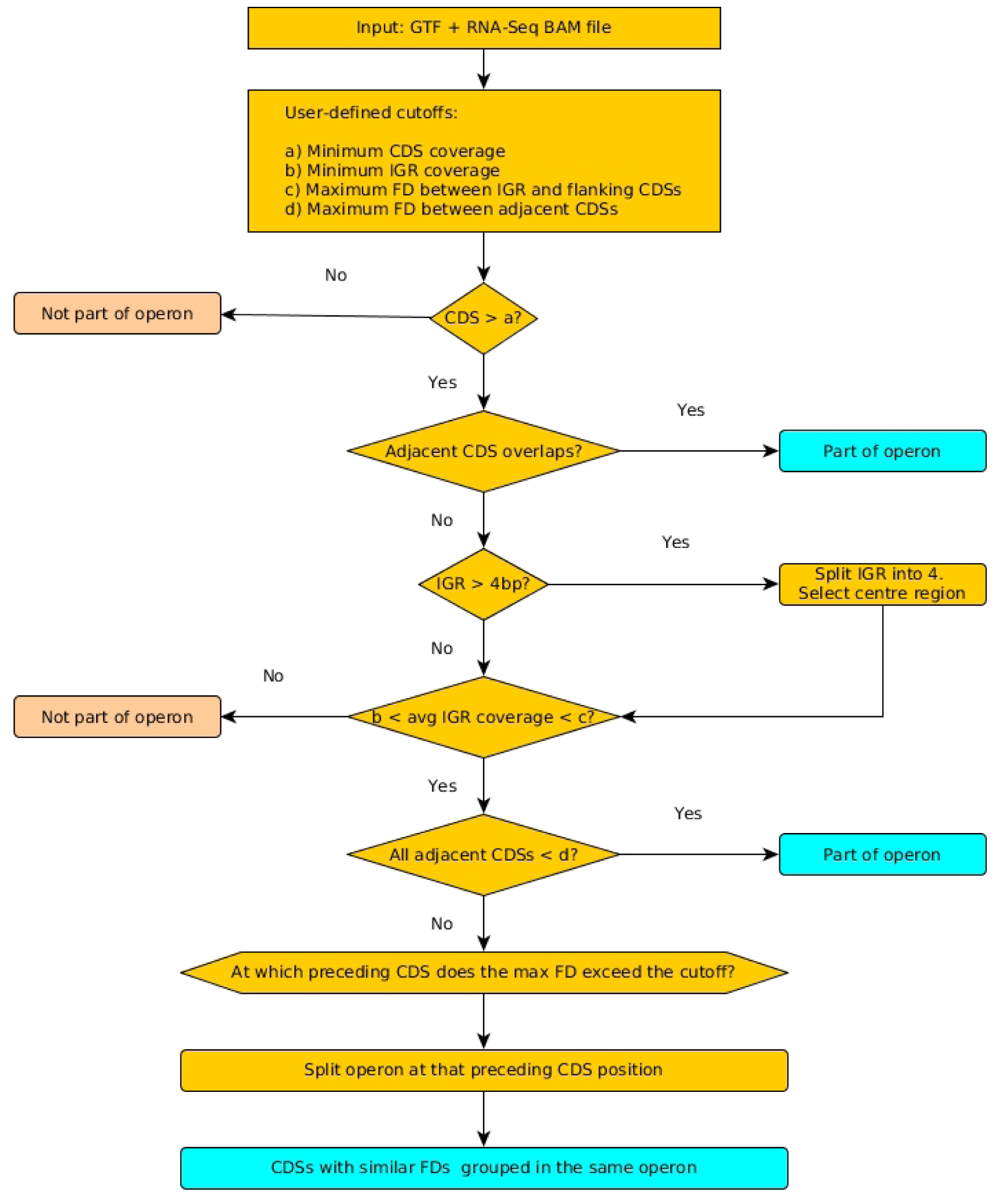
Flow diagram of COSMO’s workflow. The algorithm takes a bam file, a GTF file, four user defined cutoffs, together with some coordinate information on the genome. It then adds a CDSs to an operon if it satisfies all four conditions; the average coverage must be: ***a)*** equal to or above the CDS cutoff, ***b)*** equal to or above IGR cutoff, ***c*)** less than or equal to the maximum FD between an IGR and its flanking CDSs, and ***d)*** less than or equal to the maximum FD between adjacent CDSs.

The same process is followed with CDS 3. However, from CDS 3 and onwards, an additional rule is applied. For this fifth variable, the maximum FD is not just checked between CDS 3 and CDS 2, but the FD between CDS 3 and CDS1 must also not exceed the max-FD. If it does, then the operon is paused. However, COSMO does not automatically assume that because the coverages of CDS 1 and CDS 2 previously correlated, that they should remain an operon, and that CDS 2 should be the start of a putative new operon. Since the correlation ended when CDS 3 was compared to CDS 1, COSMO will conclude that the problem lies at CDS 1. It will therefore make a decision about how the operon should be split at CDS 1. It will evaluate whether the coverage of CDS 1 correlates better with CDS 2 and should therefore result in a bicistronic operon CDS 1 + CDS 2 or whether CDS 2 correlates better with CDS 3 and therefore become the bicistronic operon CDS 2 + CDS 3, with CDS 1 being expressed independently. This variable was a built-in feature in COSMO which could not be tested in the MLR. Although we suspected that this is likely to have a significant effect on the outcome variable.

Lastly, COSMO accounts for the circular chromosome of *Mtb*, so the first and last CDSs of the genome, may also form an operon. COSMO predicts strand specific, condition-dependent operons and outputs a CSV file. The output file contains the operon name and coordinates, the operon length and the average coverage of the operon, as well as the name and coverages of each individual gene and IGR within the operon.

#### 5.3 Algorithm validation

Although previous cutoffs were statistically validated, we wanted to confirm these cutoffs by testing a range of **actual cutoff values**. We also wanted to ascertain if having all these options made a difference in terms of the total correct operon predictions. The algorithm was therefore run on nine Beijing lineage isolates, using permutations of the following cutoffs (python and bash scripts):

a) min CDS cutoffs: [1, 2, 3, 4, 5, 6, 7, 8, 9, 10, 15, 20]
b) min IGR cutoffs: [1, 2, 3, 4, 5, 6, 7, 8, 9, 10, 15, 20]
c) max FD between adjacent CDSs: [5, 6, 7, 8, 9, 10, 15, 20]
d) max FD between IGR and flanking CDSs: [5, 6, 7, 8, 9, 10, 15, 20]

This produced 9216 files per bam file/isolate. A python script was used to compute the percentage of **full-length** operons from our EVOs list that were correctly predicted, or the proportion of TPs. We also calculated the percentage of false positives (FPs) and false negatives (FNs).

#### 5.3.1 Multiple Linear Regression Analysis

We used the data from these 9216 files to perform a MLR analysis in R Studio to observe whether each of the four parameters (independent variables), namely: min-CDS, min-IGR, FD CDSs, and FD CDS-IGR, had a statistically significant impact on the **outcome variable**. This was additionally used to verify **default** cutoff values. The outcome variable was the percentage of correctly predicted full-length operons, per combination of the different cutoff values. We performed a backwards stepwise analysis to remove any predictors that had no impact on the outcome. We also used the ‘*rpart’* [45] and ‘*rattle’* packages [46] in R Studio to draw a pruned decision tree to confirm the default parameters in COSMO, for users who wish to run COSMO on default settings. Training and test sets were split into 70% and 30%, respectively. We reported the significant predictors, as per their t-statistic p-values, R^2^, the MAE and the RMSE. R^2^ is the proportion of variation in the outcome variable that can be explained by the independent variables. The mean absolute error is an error statistic that averages the distances between each pair of actual versus observed data points (residuals) [47]. The RMSE, gives us the standard deviations of the residuals from a model. This is often argued to be the more meaningful measure of a model’s fit than the R^2^ metric [21]. The usual practice is to choose the model which has a lower accuracy measure among alternative models [47].

#### 5.3.2 Comparison to existing algorithms

Finally, the performance of COSMO was tested against other existing algorithms that use RNA-seq data as input. Initially we wanted to use DOOR 2.0, because DOOR was previously ranked as the best performing algorithm among 14 operon predicting algorithms (Mao et al., 2009). Unfortunately, it had become obsolete at the time of our testing. We therefore compared our results to Rockhopper and REMap. REMap was chosen because the algorithm’s approach was similar to ours and tailor-made for the Mtb genome. Further encouragement was also due to REMap’s performance which fared on par with DOOR 2.0 [1]. Rockhopper was our second comparator, because it was previously shown to outperform DOOR 2.0 [48].

REMap published that an expression level of 10x was able to yield the best results. The algorithm however had a default value of 20x. Therefore, both parameters were used to predict operons using our datasets. Rockhopper does not allow user-defined expression level cutoffs.

The total number of correctly predicted operons (TPs), as well as the total number of FPs and FNs, were calculated for each predictor. The algorithms were evaluated with the performance metrics: precision/positive predictive value (PPV), recall (sensitivity), and F1 score, where:

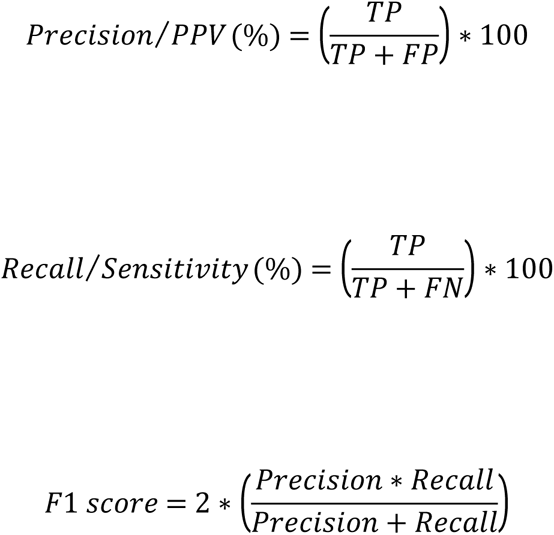

In the absence of a verified true negative operon set, we could not calculate the specificities, accuracy scores or ROC curves. Secondly, the actual number of operons compared to single genes most likely results in an unbalanced dataset. In these two scenarios, the F1 score has been proven to be a better metric than the accuracy score to evaluate algorithm performance [49]. Similarly, when datasets are unbalanced, precision and recall were demonstrated as better evaluators for a model’s classification performance; and precision-recall curves were more useful and robust than the ROC curves.

## Acknowledgements

We are sincerely grateful to South African Medical Research Council Bioinformatics Unit, South African National Bioinformatics Institute, University of the Western Cape, South Africa (SAMRC/SANBI) for assistance with funding for this project. We also wish to thank Dr Jean-Baka Domelevo Entfellner (BecA-ILRI) and Dr Sunday Vodah (previous SANBI post-doc) for assisting with some of the statistics and the mathematics, respectively. Also thank you to the SANBI staff: Thoba Lose, Zipho Mashologu, Quinton Coert, Eugene de Beste, who at times had to assist with the scripting.

## 8 Supporting Information

**S1 Table 1:**
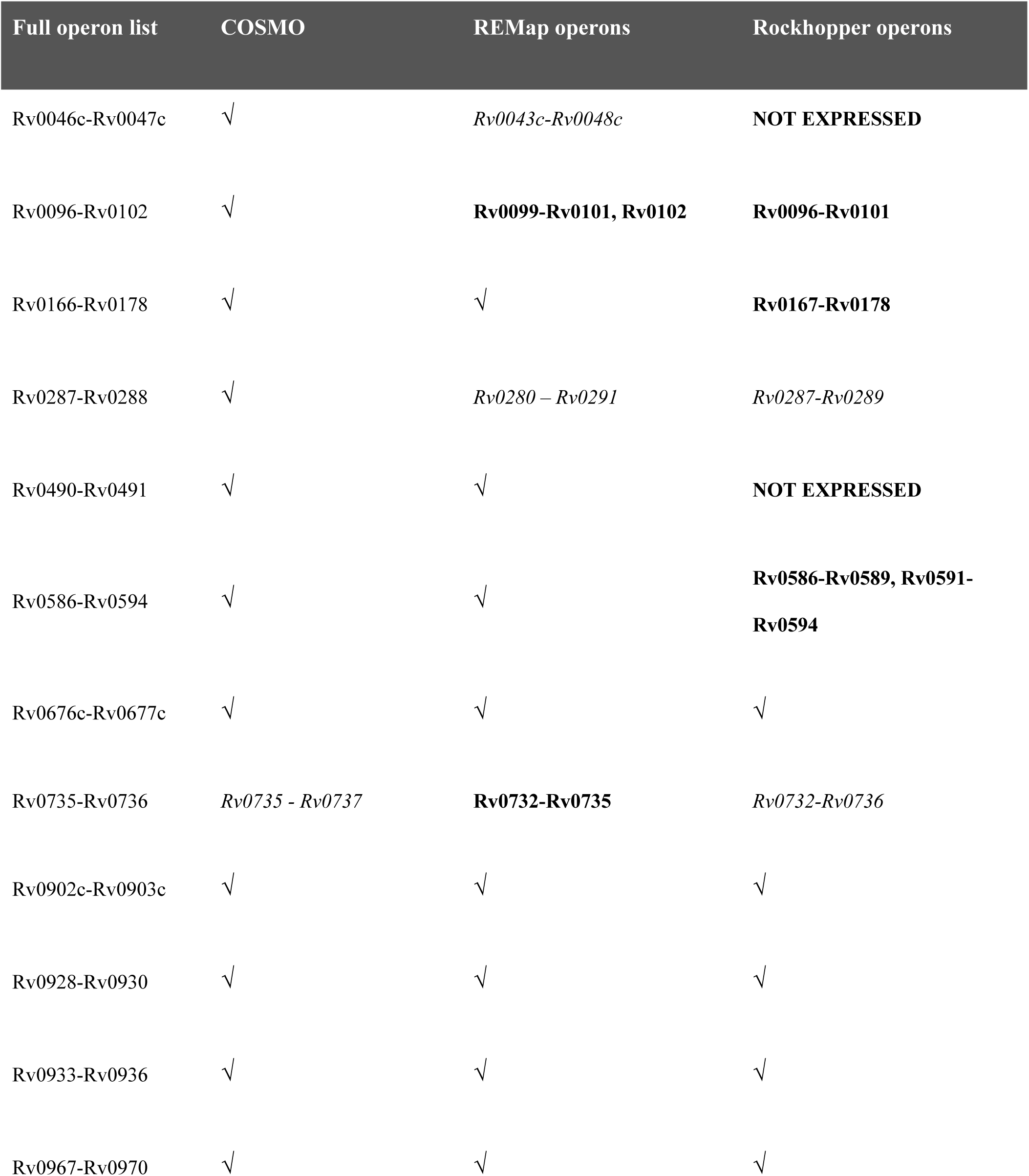
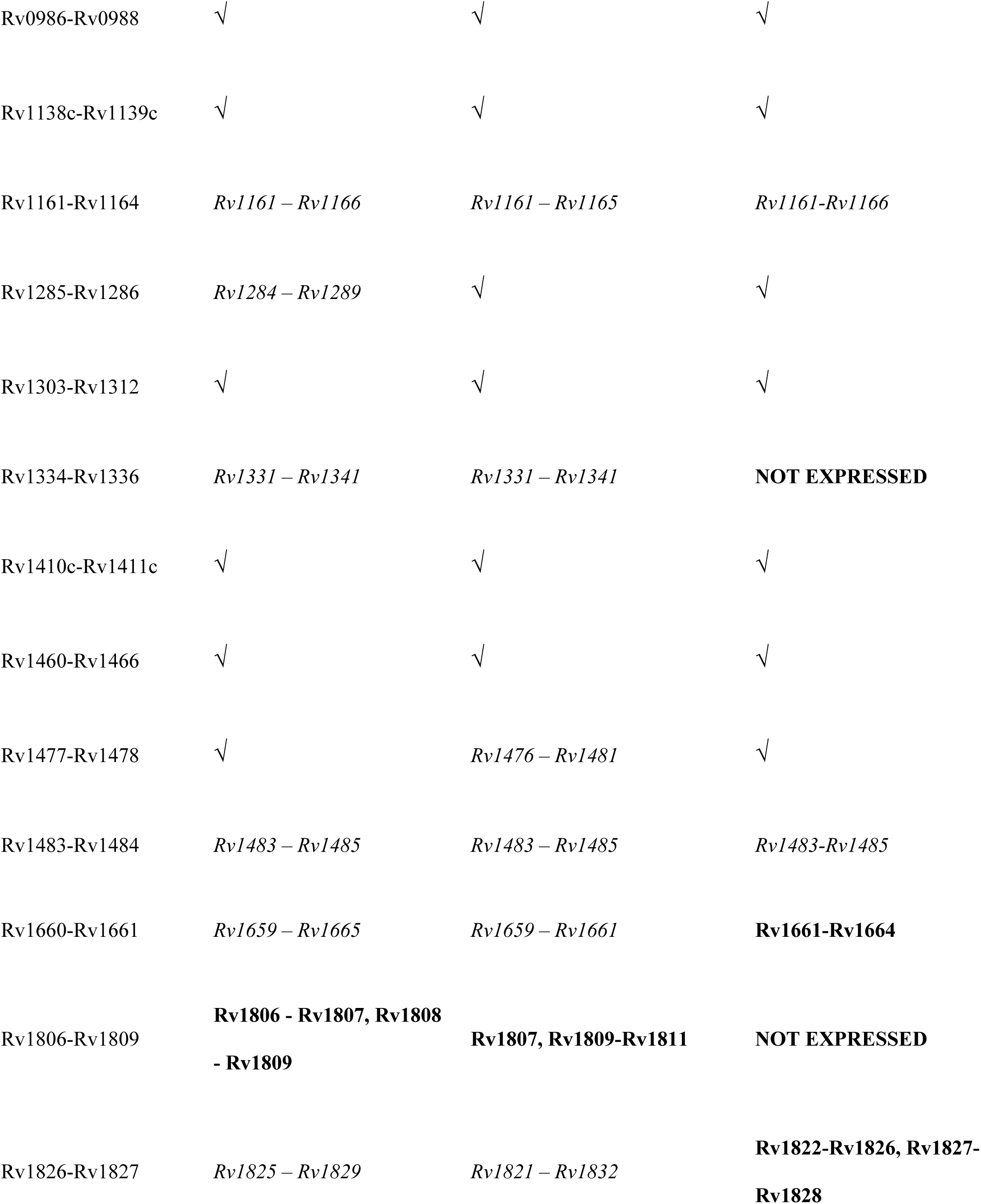
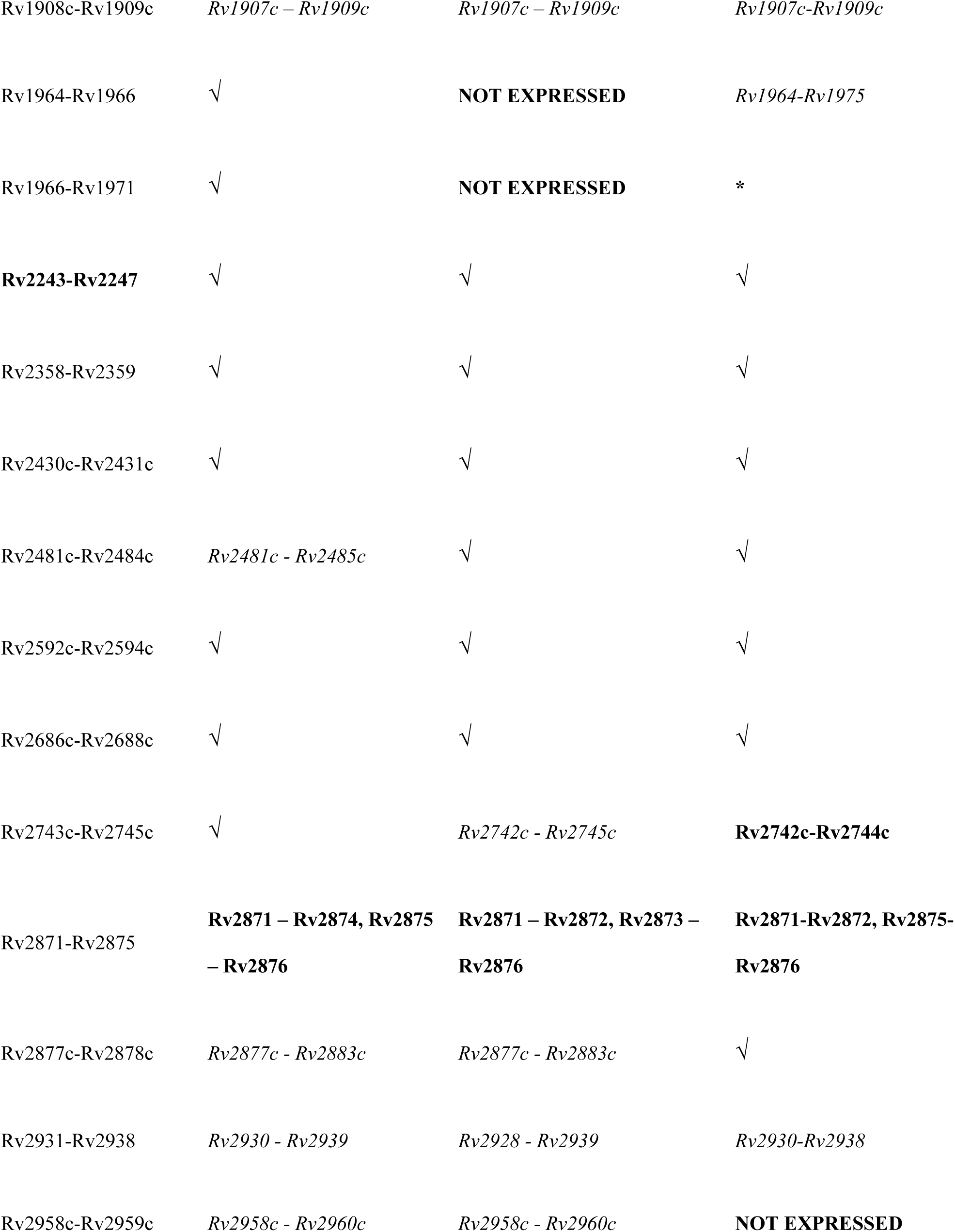
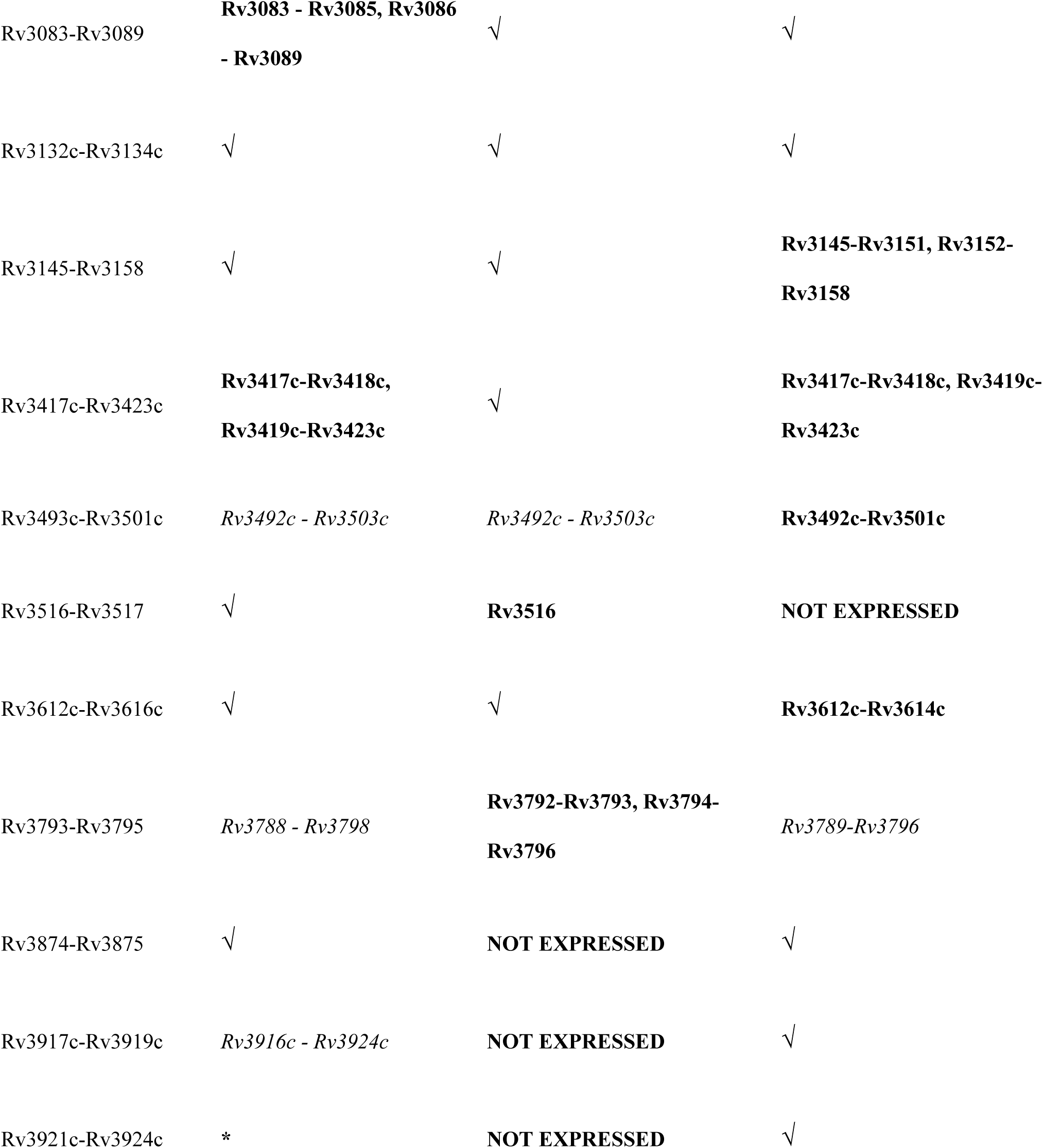
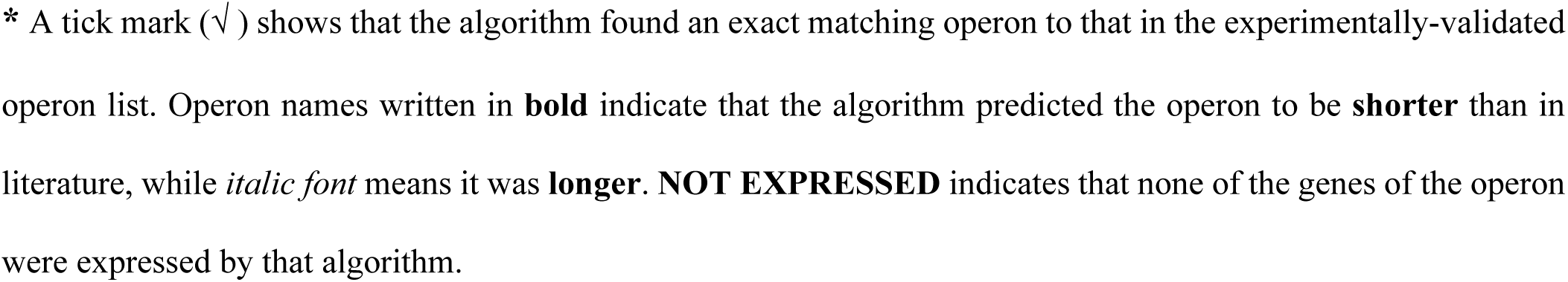
Correctly predicted operons made by the three operon algorithms compared to all the experimentally validated operons.

| Full operon list | COSMO | REMap operons | Rockhopper operons |
| --- | --- | --- | --- |
| Rv0046c-Rv0047c | ✓ | *Rv0043c-Rv0048c* | **NOT EXPRESSED** |
| Rv0096-Rv0102 | ✓ | **Rv0099-Rv0101, Rv0102** | **Rv0096-Rv0101** |
| Rv0166-Rv0178 | ✓ | ✓ | **Rv0167-Rv0178** |
| Rv0287-Rv0288 | ✓ | *Rv0280 – Rv0291* | *Rv0287-Rv0289* |
| Rv0490-Rv0491 | ✓ | ✓ | **NOT EXPRESSED** |
| Rv0586-Rv0594 | ✓ | ✓ | **Rv0586-Rv0589, Rv0591-**<br>**Rv0594** |
| Rv0676c-Rv0677c | ✓ | ✓ | ✓ |
| Rv0735-Rv0736 | *Rv0735 - Rv0737* | **Rv0732-Rv0735** | *Rv0732-Rv0736* |
| Rv0902c-Rv0903c | ✓ | ✓ | ✓ |
| Rv0928-Rv0930 | ✓ | ✓ | ✓ |
| Rv0933-Rv0936 | ✓ | ✓ | ✓ |
| Rv0967-Rv0970 | ✓ | ✓ | ✓ |
| Rv0986-Rv0988 | ✓ | ✓ | ✓ |
| Rv1138c-Rv1139c | ✓ | ✓ | ✓ |
| Rv1161-Rv1164 | *Rv1161 – Rv1166* | *Rv1161 – Rv1165* | *Rv1161-Rv1166* |
| Rv1285-Rv1286 | *Rv1284 – Rv1289* | ✓ | ✓ |
| Rv1303-Rv1312 | ✓ | ✓ | ✓ |
| Rv1334-Rv1336 | *Rv1331 – Rv1341* | *Rv1331 – Rv1341* | **NOT EXPRESSED** |
| Rv1410c-Rv1411c | ✓ | ✓ | ✓ |
| Rv1460-Rv1466 | ✓ | ✓ | ✓ |
| Rv1477-Rv1478 | ✓ | *Rv1476 – Rv1481* | ✓ |
| Rv1483-Rv1484 | *Rv1483 – Rv1485* | *Rv1483 – Rv1485* | *Rv1483-Rv1485* |
| Rv1660-Rv1661 | *Rv1659 – Rv1665* | *Rv1659 – Rv1661* | **Rv1661-Rv1664** |
| Rv1806-Rv1809 | **Rv1806 - Rv1807, Rv1808**<br>**- Rv1809** | **Rv1807, Rv1809-Rv1811** | **NOT EXPRESSED** |
| Rv1826-Rv1827 | *Rv1825 – Rv1829* | *Rv1821 – Rv1832* | **Rv1822-Rv1826, Rv1827-**<br>**Rv1828** |
| Rv1908c-Rv1909c | *Rv1907c – Rv1909c* | *Rv1907c – Rv1909c* | *Rv1907c-Rv1909c* |
| Rv1964-Rv1966 | √ | **NOT EXPRESSED** | *Rv1964-Rv1975* |
| Rv1966-Rv1971 | √ | **NOT EXPRESSED** | * |
| **Rv2243-Rv2247** | √ | √ | √ |
| Rv2358-Rv2359 | √ | √ | √ |
| Rv2430c-Rv2431c | √ | √ | √ |
| Rv2481c-Rv2484c | *Rv2481c - Rv2485c* | √ | √ |
| Rv2592c-Rv2594c | √ | √ | √ |
| Rv2686c-Rv2688c | √ | √ | √ |
| Rv2743c-Rv2745c | √ | *Rv2742c - Rv2745c* | **Rv2742c-Rv2744c** |
| Rv2871-Rv2875 | **Rv2871 – Rv2874, Rv2875**<br>**– Rv2876** | **Rv2871 – Rv2872, Rv2873 –**<br>**Rv2876** | **Rv2871-Rv2872, Rv2875-**<br>**Rv2876** |
| Rv2877c-Rv2878c | *Rv2877c - Rv2883c* | *Rv2877c - Rv2883c* | √ |
| Rv2931-Rv2938 | *Rv2930 - Rv2939* | *Rv2928 - Rv2939* | *Rv2930-Rv2938* |
| Rv2958c-Rv2959c | *Rv2958c - Rv2960c* | *Rv2958c - Rv2960c* | **NOT EXPRESSED** |
| Rv3083-Rv3089 | **Rv3083 - Rv3085, Rv3086**<br>**- Rv3089** | √ | √ |
| Rv3132c-Rv3134c | √ | √ | √ |
| Rv3145-Rv3158 | √ | √ | **Rv3145-Rv3151, Rv3152-**<br>**Rv3158** |
| Rv3417c-Rv3423c | **Rv3417c-Rv3418c,**<br>**Rv3419c-Rv3423c** | √ | **Rv3417c-Rv3418c, Rv3419c-**<br>**Rv3423c** |
| Rv3493c-Rv3501c | *Rv3492c - Rv3503c* | *Rv3492c - Rv3503c* | **Rv3492c-Rv3501c** |
| Rv3516-Rv3517 | √ | **Rv3516** | **NOT EXPRESSED** |
| Rv3612c-Rv3616c | √ | √ | **Rv3612c-Rv3614c** |
| Rv3793-Rv3795 | *Rv3788 - Rv3798* | **Rv3792-Rv3793, Rv3794-**<br>**Rv3796** | *Rv3789-Rv3796* |
| Rv3874-Rv3875 | √ | **NOT EXPRESSED** | √ |
| Rv3917c-Rv3919c | *Rv3916c - Rv3924c* | **NOT EXPRESSED** | √ |
| Rv3921c-Rv3924c | * | **NOT EXPRESSED** | √ |

\* A tick mark (✓) shows that the algorithm found an exact matching operon to that in the experimentally-validated operon list. Operon names written in **bold** indicate that the algorithm predicted the operon to be **shorter** than in literature, while *italic font* means it was **longer**. **NOT EXPRESSED** indicates that none of the genes of the operon were expressed by that algorithm.

**S2 Fig 1:**
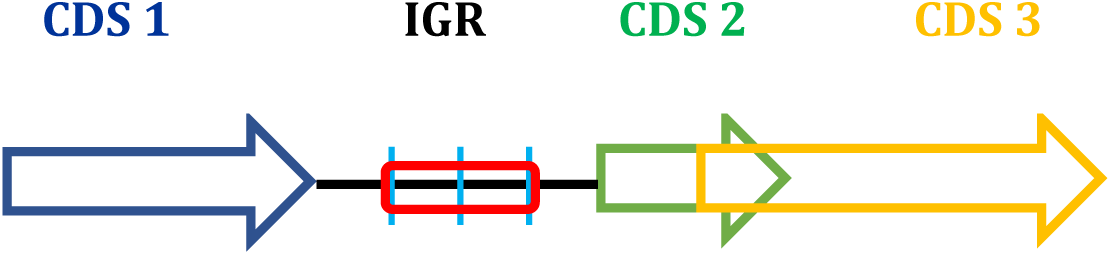
Illustration of the central region of an intergenic region relative to flanking CDSs. Two CDSs may be separated by an intergenic region (IGR) such as depicted with the two adjacent CDSs: CDS 1 and CDS 2. CDSs may also overlap such as the case with CDS 2 and CDS 3. In the case where they do not overlap, we calculate the FD between an IGR and each flanking CDS. However, we do not consider the entire IGR – which is the entire black line extending from where CDS 1 ends, to where CDS 2 begins, but only the centre of the IGR (red block). The entire IGR is only used when the IGR is <= 4bp.

## References

1. Pelly S, Winglee K, Xia FF, Stevens RL, Bishai WR, Lamichhane G. REMap: Operon Map of M. tuberculosis. Tuberc Edinb Scotl [Internet]. 2016 [cited 2019 Oct 19];99:70–80. Available from: https://www.ncbi.nlm.nih.gov/pmc/articles/PMC4967370/

2. Price MN, Arkin AP, Alm EJ. The Life-Cycle of Operons. PLOS Genet [Internet]. 2006 [cited 2019 Oct 19];2:e96. Available from: https://journals.plos.org/plosgenetics/article?id=10.1371/journal.pgen.0020096

3. Osbourn AE, Field B. Operons. Cell Mol Life Sci [Internet]. 2009 [cited 2019 Oct 19];66:3755–75. Available from: https://doi.org/10.1007/s00018-009-0114-3

4. Güell M, van Noort V, Yus E, Chen W-H, Leigh-Bell J, Michalodimitrakis K, et al. Transcriptome complexity in a genome-reduced bacterium. Science. 2009;326:1268–71.

5. Zaidi SSA, Zhang X. Computational operon prediction in whole-genomes and metagenomes. Brief Funct Genomics [Internet]. Oxford Academic; 2017 [cited 2020 Apr 18];16:181–93. Available from: https://academic.oup.com/bfg/article/16/4/181/2555398

6. Zhao S, Prenger K, Smith L. Stormbow: A Cloud-Based Tool for Reads Mapping and Expression Quantification in Large-Scale RNA-Seq Studies. ISRN Bioinforma [Internet]. 2013 [cited 2021 Sep 13];2013:481545. Available from: https://www.ncbi.nlm.nih.gov/pmc/articles/PMC4393068/

7. Rao MS, Van Vleet TR, Ciurlionis R, Buck WR, Mittelstadt SW, Blomme EAG, et al. Comparison of RNA-Seq and Microarray Gene Expression Platforms for the Toxicogenomic Evaluation of Liver From Short-Term Rat Toxicity Studies. Front Genet [Internet]. Frontiers; 2019 [cited 2021 Jun 19];9. Available from: https://www.frontiersin.org/articles/10.3389/fgene.2018.00636/full

8. Tjaden B. A computational system for identifying operons based on RNA-seq data. Methods. 2020;176:62–70.

9. Cao H, Ma Q, Chen X, Xu Y. DOOR: a prokaryotic operon database for genome analyses and functional inference. Brief Bioinform. 2019;20:1568–77.

10. Che D, Li G, Mao F, Wu H, Xu Y. Detecting uber-operons in prokaryotic genomes. Nucleic Acids Res [Internet]. 2006 [cited 2019 Dec 11];34:2418–27. Available from: https://www.ncbi.nlm.nih.gov/pmc/articles/PMC1458513/

11. Bergman NH, Passalacqua KD, Hanna PC, Qin ZS. Operon Prediction for Sequenced Bacterial Genomes without Experimental Information. Appl Environ Microbiol [Internet]. 2007 [cited 2019 Dec 11];73:846–54. Available from: https://aem.asm.org/content/73/3/846

12. Bashyam MD, Kaushal D, Dasgupta SK, Tyagi AK. A study of mycobacterial transcriptional apparatus: identification of novel features in promoter elements. J Bacteriol. 1996;178:4847–53.

13. Cortes T, Schubert OT, Rose G, Arnvig KB, Comas I, Aebersold R, et al. Genome-wide mapping of transcriptional start sites defines an extensive leaderless transcriptome in Mycobacterium tuberculosis. Cell Rep. 2013;5:1121–31.

14. Chuang L-Y, Chang H-W, Tsai J-H, Yang C-H. Features for computational operon prediction in prokaryotes. Brief Funct Genomics [Internet]. 2012 [cited 2020 Jan 30];11:291–9. Available from: https://academic.oup.com/bfg/article/11/4/291/200556

15. Fortino V, Smolander O-P, Auvinen P, Tagliaferri R, Greco D. Transcriptome dynamics-based operon prediction in prokaryotes. BMC Bioinformatics [Internet]. 2014 [cited 2019 Dec 11];15:145. Available from: https://doi.org/10.1186/1471-2105-15-145

16. Salgado H, Moreno-Hagelsieb G, Smith TF, Collado-Vides J. Operons in Escherichia coli: Genomic analyses and predictions. Proc Natl Acad Sci [Internet]. 2000 [cited 2020 Jan 29];97:6652–7. Available from: https://www.pnas.org/content/97/12/6652

17. Taboada B, Estrada K, Ciria R, Merino E. Operon-mapper: a web server for precise operon identification in bacterial and archaeal genomes. Bioinformatics [Internet]. 2018 [cited 2019 Dec 11];34:4118–20. Available from: https://academic.oup.com/bioinformatics/article/34/23/4118/5040321

18. Aravindhan V, Christy AJ, Roy S, Ajitkumar P, Narayanan PR, Narayanan S. Mycobacterium tuberculosis groE promoter controls the expression of the bicistronic groESL1 operon and shows differential regulation under stress conditions. FEMS Microbiol Lett [Internet]. Oxford Academic; 2009 [cited 2020 Nov 10];292:42–9. Available from: https://academic.oup.com/femsle/article/292/1/42/490157

19. Bhat AH, Pathak D, Rao A. The alr-groEL1 operon in Mycobacterium tuberculosis : an interplay of multiple regulatory elements. Sci Rep [Internet]. Nature Publishing Group; 2017 [cited 2020 Nov 10];7:43772. Available from: https://www.nature.com/articles/srep43772

20. Stewart GR, Wernisch L, Stabler R, Mangan JA, Hinds J, Laing KG, et al. Dissection of the heat-shock response in Mycobacterium tuberculosis using mutants and microarraysaaA list of the 100 ORFs most highly induced by heat shock is provided as supplementary data with the online version of this paper (http://mic.sgmjournals.org). Microbiology, [Internet]. Microbiology Society,; 2002 [cited 2020 Nov 10];148:3129–38. Available from: https://www.microbiologyresearch.org/content/journal/micro/10.1099/00221287-148-10-3129

21. Alexander DLJ, Tropsha A, Winkler DA. Beware of R2: simple, unambiguous assessment of the prediction accuracy of QSAR and QSPR models. J Chem Inf Model [Internet]. 2015 [cited 2021 Oct 6];55:1316–22. Available from: https://www.ncbi.nlm.nih.gov/pmc/articles/PMC4530125/

22. Li J. Assessing the accuracy of predictive models for numerical data: Not r nor r2, why not? Then what? PLOS ONE [Internet]. Public Library of Science; 2017 [cited 2021 Oct 6];12:e0183250. Available from: https://journals.plos.org/plosone/article?id=10.1371/journal.pone.0183250

23. Bockhorst J, Craven M, Page D, Shavlik J, Glasner J. A Bayesian network approach to operon prediction. Bioinformatics [Internet]. 2003 [cited 2017 Aug 25];19:1227–35. Available from: https://academic.oup.com/bioinformatics/article/19/10/1227/184417/A-Bayesian-network-approach-to-operon-prediction

24. Laing E, Sidhu K, Hubbard SJ. Predicted transcription factor binding sites as predictors of operons in Escherichia coli and Streptomyces coelicolor. BMC Genomics. 2008;9:79.

25. McClure R, Balasubramanian D, Sun Y, Bobrovskyy M, Sumby P, Genco CA, et al. Computational analysis of bacterial RNA-Seq data. Nucleic Acids Res [Internet]. 2013 [cited 2019 Oct 23];41:e140. Available from: https://www.ncbi.nlm.nih.gov/pmc/articles/PMC3737546/

26. Zheng Y, Szustakowski JD, Fortnow L, Roberts RJ, Kasif S. Computational identification of operons in microbial genomes. Genome Res. 2002;12:1221–30.

27. Hunter JD. Matplotlib: A 2D Graphics Environment. Comput Sci Eng [Internet]. 2007 [cited 2022 Mar 23];9:90–5. Available from: https://doi.org/10.1109/MCSE.2007.55

28. Kluyver T, Ragan-Kelley B, Pérez F, Granger B, Bussonnier M, Frederic J, et al. Jupyter Notebooks – a publishing format for reproducible computational workflows. In: Loizides F, Scmidt B, editors. IOS Press; 2016 [cited 2022 Apr 27]. p. 87–90. Available from: https://eprints.soton.ac.uk/403913/

29. Namouchi A, Gómez-Muñoz M, Frye SA, Moen LV, Rognes T, Tønjum T, et al. The Mycobacterium tuberculosis transcriptional landscape under genotoxic stress. BMC Genomics [Internet]. 2016 [cited 2017 Feb 3];17:791. Available from: http://dx.doi.org/10.1186/s12864-016-3132-1

30. World Health Organization. Global tuberculosis report 2019 [Internet]. 2019. Available from: https://apps.who.int/iris/bitstream/handle/10665/329368/9789241565714-eng.pdf?ua=1

31. Coll F, Phelan J, Hill-Cawthorne GA, Nair MB, Mallard K, Ali S, et al. Genome-wide analysis of multi- and extensively drug-resistant Mycobacterium tuberculosis. Nat Genet [Internet]. Nature Publishing Group; 2018 [cited 2020 Apr 17];50:307–16. Available from: https://www.nature.com/articles/s41588-017-0029-0

32. Hoagland DT, Liu J, Lee RB, Lee RE. New agents for the treatment of drug-resistant Mycobacterium tuberculosis. Adv Drug Deliv Rev [Internet]. 2016 [cited 2020 Apr 17];102:55–72. Available from: http://www.sciencedirect.com/science/article/pii/S0169409X16301363

33. Phelan JE, O’Sullivan DM, Machado D, Ramos J, Oppong YEA, Campino S, et al. Integrating informatics tools and portable sequencing technology for rapid detection of resistance to anti-tuberculous drugs. Genome Med [Internet]. 2019 [cited 2022 Apr 27];11:41. Available from: https://doi.org/10.1186/s13073-019-0650-x

34. Jalili V, Afgan E, Gu Q, Clements D, Blankenberg D, Goecks J, et al. The Galaxy platform for accessible, reproducible and collaborative biomedical analyses: 2020 update. Nucleic Acids Res. 2020;48:W395–402.

35. Andrews S. Babraham Bioinformatics - FastQC A Quality Control tool for High Throughput Sequence Data [Internet]. [cited 2022 Apr 27]. Available from: https://www.bioinformatics.babraham.ac.uk/projects/fastqc/

36. Bolger AM, Lohse M, Usadel B. Trimmomatic: a flexible trimmer for Illumina sequence data. Bioinformatics [Internet]. 2014 [cited 2022 Mar 23];30:2114–20. Available from: https://doi.org/10.1093/bioinformatics/btu170

37. Li H, Durbin R. Fast and accurate short read alignment with Burrows–Wheeler transform. Bioinformatics [Internet]. 2009 [cited 2022 Mar 23];25:1754–60. Available from: https://www.ncbi.nlm.nih.gov/pmc/articles/PMC2705234/

38. Li H, Handsaker B, Wysoker A, Fennell T, Ruan J, Homer N, et al. The Sequence Alignment/Map format and SAMtools. Bioinformatics [Internet]. 2009 [cited 2022 Mar 23];25:2078–9. Available from: https://www.ncbi.nlm.nih.gov/pmc/articles/PMC2723002/

39. Wang L, Wang S, Li W. RSeQC: quality control of RNA-seq experiments. Bioinformatics [Internet]. 2012 [cited 2022 Mar 23];28:2184–5. Available from: https://doi.org/10.1093/bioinformatics/bts356

40. Tjaden B, Saxena RM, Stolyar S, Haynor DR, Kolker E, Rosenow C. Transcriptome analysis of Escherichia coli using high-density oligonucleotide probe arrays. Nucleic Acids Res [Internet]. 2002 [cited 2021 Sep 4];30:3732–8. Available from: https://www.ncbi.nlm.nih.gov/pmc/articles/PMC137427/

41. Okuda S, Kawashima S, Kobayashi K, Ogasawara N, Kanehisa M, Goto S. Characterization of relationships between transcriptional units and operon structures in Bacillus subtilis and Escherichia coli. BMC Genomics [Internet]. 2007 [cited 2021 Sep 3];8:48. Available from: https://www.ncbi.nlm.nih.gov/pmc/articles/PMC1808063/

42. Arnvig KB, Comas I, Thomson NR, Houghton J, Boshoff HI, Croucher NJ, et al. Sequence-Based Analysis Uncovers an Abundance of Non-Coding RNA in the Total Transcriptome of Mycobacterium tuberculosis. PLOS Pathog [Internet]. 2011 [cited 2016 Aug 10];7:e1002342. Available from: http://journals.plos.org/plospathogens/article?id=10.1371/journal.ppat.1002342

43. Sedlyarova N, Shamovsky I, Bharati BK, Epshtein V, Chen J, Gottesman S, et al. sRNA-mediated control of transcription termination in E. coli. Cell [Internet]. 2016 [cited 2022 Mar 25];167:111–121.e13. Available from: https://www.ncbi.nlm.nih.gov/pmc/articles/PMC5040353/

44. R Core Team. R: A Language and Environment for Statistical Computing [Internet]. 2021. Available from: https://www.R-project.org

45. Therneau T, Atkinson E. An introduction to recursive partitioning using the RPART routines [Internet]. 2022. Available from: https://mran.microsoft.com/web/packages/rpart/rpart.pdf

46. Williams G. Data Mining with Rattle and R: The Art of Excavating Data for Knowledge Discovery. Springer Science & Business Media; 2011.

47. Boiroju N. A Bootstrap Test for Equality of Mean Absolute Errors. ARPN J Eng Appl Sci. 2011;

48. Tjaden B. A computational system for identifying operons based on RNA-seq data. Methods [Internet]. 2019 [cited 2019 Oct 23]; Available from: http://www.sciencedirect.com/science/article/pii/S1046202318303426

49. DeVries Z, Locke E, Hoda M, Moravek D, Phan K, Stratton A, et al. Using a national surgical database to predict complications following posterior lumbar surgery and comparing the area under the curve and F1-score for the assessment of prognostic capability. Spine J Off J North Am Spine Soc. 2021;21:1135–42.

